# A variant centric analysis of allele sharing in dogs and wolves

**DOI:** 10.1101/2024.08.08.607131

**Authors:** Matthew W. Funk, Jeffrey M. Kidd

**Affiliations:** Department of Computational Medicine & Bioinformatics, University of Michigan, Ann Arbor, MI; Department of Human Genetics, University of Michigan, Ann Arbor, MI

**Keywords:** Canine genetics, allele sharing, population structure

## Abstract

Canines are an important model system for genetics and evolution. Recent advances in sequencing technologies have enabled the creation of large databases of genetic variation in canines, but analysis of allele sharing among canine groups has been limited. We applied GeoVar, an approach originally developed to study the sharing of single nucleotide polymorphisms across human populations, to assess the sharing of genetic variation among groups of wolves, village dogs, and breed dogs. Our analysis shows that wolves differ from each other at an average of approximately 2.3 million sites while dogs from the same breed differ at nearly 1 million sites. We find that 22% of variants are common across wolves, village dogs, and breed dogs, that ∼16% of variable sites are common across breed dogs, and that nearly half of the differences between two dogs of different breeds are due to sites that are common in all clades. These analyses represent a succinct summary of allele sharing across canines and illustrate the effects of canine history on the apportionment of genetic variation.

## 1. Introduction

Domestic dogs (*Canis lupus familiaris)* are a powerful model system for genetics and evolutionary biology. Genetic evidence from modern and ancient samples indicate that dogs were domesticated from a now-extinct lineage of wolves 20,000 to 30,000 years ago somewhere in Eurasia [1-3]. Global surveys of dog genetic diversity show a clear separation between samples of western Eurasian and eastern Eurasian origin [4]. Since domestication, dogs have experienced a complex demographic history including waves of global migration that parallel the movement of human populations [3]. These demographic events include bottlenecks, expansions, and periods of interbreeding and population replacement that have left distinct signatures in the patterns of canine genetic diversity [5,6]. Modern dog breeds, which are genetically closed populations that are bred towards specified phenotypes, are a relatively recent development with origins within the past 400 years, with most breeds originating during the 19th century [7]. The unique genetic structure of modern dog breeds facilitates the identification of alleles affecting disease susceptibility, morphology and behavior [8].

The severe bottlenecks and sustained small effective population sizes associated with breed formation have had profound effects on canine genomes including an increased load of deleterious alleles [6], increased presence of recessively inherited disorders [9], and an associated decrease in overall fitness [10-12]. Despite their large phenotypic diversity and popularity as companion animals, most canine genetic variation is not found among breed dogs. Rather, most genetic diversity is found in populations of dogs that live as semi-feral human commensals around the world [13,14]. These populations, known as village dogs or street dogs, better represent the state of dogs throughout their long history prior to the formation of modern breeds. Importantly, genetic studies have confirmed that village dogs represent distinct populations that have retained the genetic diversity lost during the formation of breeds and are not simply mixtures of breed dogs that have “escaped”[15]. Thus, breed dogs, village dogs, and wolves reflect distinct aspects of canine history.

The relevance of dogs to human disease studies has led to the growth of a robust canine genetics research community. This research community has developed valuable resources including a well-annotated reference genome [16,17], detailed phenotype information coupled with databases of known genetic variation that enable efficient genome wide trait mapping [18], and publicly available collections of samples with short-read whole genome sequence data [19-21]. This strong research foundation has been expanded by the recent availability of multiple high-quality reference genomes derived from long-read sequencing technologies [22-29]. Recently, the Dog10K consortium released whole genome sequencing data from a diverse collection of nearly 2,000 samples including wolves, village dogs, and breed dogs [21]. This collection offers an unbiased view of canine genome variation and is a valuable resource for trait mapping and evolutionary studies. Importantly, this sequence-based resource represents a genome-wide perspective on patterns of genetic variation in canines without the biases associated with genotyping arrays [30,31]. However, effectively visualizing the sharing of genetic variation among multiple sample categories in this large collection remains a challenge.

Visualizing the large genetic data sets generated from geographically diverse human populations has also been a challenge. In 2020 Biddanda, Rice, and Novembre developed a technique, known as GeoVar, to summarize the sharing patterns of alleles of different frequencies [32]. The GeoVar approach bins alleles into three categories based on their frequency in a population: unobserved, coded as *U*; rare, coded as *R*; and common, coded as *C*. The joint distribution of frequencies across populations is then conveyed using the encoding for each variant. For example, a variant that is common in each of four populations will be coded as *‘CCCC’*, while a variant that is rare in the first population and unobserved in the other three will be coded as *‘RUUU’*. The overall pattern of allele frequencies can then be depicted as the relative frequency of each encoding, which is represented as a GeoVar plot [32]. Essentially, this scheme represents a discretization of the multi-population site frequency spectrum and allows for a greater understanding of how rare and common variation are distributed among various groups. Application of this approach to human data revealed that the majority of variants are rare in one geographic location and unobserved elsewhere, variants that are common in one region are likely to also be found globally, most of the differences between two individuals are due to globally common alleles, and genotyping arrays are heavily biased toward globally common alleles.

In this study, we analyze genome-wide single nucleotide polymorphism (SNP) data from the Dog10K project to investigate patterns of allele sharing among canines. First, we estimate the average number of differences found at accessible SNP positions between samples, confirming that wolves show the greatest amount of genetic variation. We then apply the GeoVar approach to breed dogs, village dogs, and wolves, as well as to smaller sub-groupings among breed and village dogs. We find that globally common alleles are more frequent in canines as compared to similarly processed human samples, that ∼71% of the sites that differ between two breed dogs or two village dogs are common throughout canines, and that canine genotyping arrays are strongly biased toward common alleles.

## 2. Material and Methods

### 2.1 Canine variant data processing

Single nucleotide polymorphism (SNP) data was obtained from the Dog10K consortium based on alignment of Illumina sequence data to the canFam4/UU_Cfam-GSD_1.0 reference assembly [21]. We utilized the final ‘strict filtering’ sample set that includes 1929 individuals consisting of 1579 breed dogs, 281 village dogs, 57 wolves, and 12 dogs with a mixed origin or that are not recognized by any international registering body. We further filtered the sites to remove variation on the sex chromosomes, non-biallelic SNPs, SNPs with missing data, and SNPs that were not variable among the analyzed samples. A total of 26,585,484 SNPs (79% of the original SNPs marked as ‘PASS’ in the VCF file) were retained.

### 2.2 SNP distance calculation

An estimate of SNP distances between a pair of samples over k biallelic SNPs was calculated as in [33] as 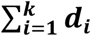 where d_i_ is 1 when the pair has opposite homozygous genotypes, ½ when one sample is homozygous and the other heterozygous, and 0 when both samples have the same genotype. This is a conservative estimate of the number of differences expected between two alleles randomly sampled from two diploid individuals as heterozygotes are assumed to be concordant. For village dogs, wolves, and within breeds, distance was calculated between all possible pairs of individuals. For the full breed dog analysis one sample from each breed was selected.

### 2.3 Canine GeoVar analysis

Allele sharing analysis was performed based on the minor allele frequency of each SNP found in the analyzed sample set using the *geovar* software as previously described [32]. Since variation discovered is sensitive to differences in sample size, analysis was performed based on a fixed number of individuals per category using a random selection of samples. For analysis of sharing among canines, 50 wolves, 50 village dogs, and 50 breed dogs were randomly selected. For analysis of village dog groups, 35 samples in each category were randomly selected. For breeds, 30 samples in each breed clade were analyzed. For each analysis, five different random samples were analyzed to assess the effect of sample selection on the overall result.

#### 2.3.1 Village dog classification

Sample groups for village dog analyses were defined based on Principal Component Analysis (PCA) and reported sample location for all village dogs in the strict filter sample set (Table S1). Prior to PCA, SNPs were filtered with PLINK version 1.9 [34] to remove SNPs with a minor allele frequency of < 5% and in high linkage disequilibrium (using PLINK *--indep-pairwise 50 10 0.1*). PCA was performed using PLINK.

#### 2.3.2 Breed dog classification

Breed dog clades were defined as in Meadows *et al*. [21]. Only clades with more than 30 individuals were retained for analysis (Table 1).

**Table 1.** Breed clades used for GeoVar analysis.

| Clade | Number of Samples | Clade | Number of Samples |
| --- | --- | --- | --- |
| Alpine | 67 | Mastiff | 97 |
| American Terriers | 31 | Pointer | 138 |
| Asian | 135 | Retriever | 36 |
| Belgian Herders | 43 | Scenthound | 222 |
| Continental Herders | 38 | Scottish Terrier | 40 |
| English Terriers | 53 | Spaniel | 62 |
| Flockguard<br>Sighthound | 145 | Spitz | 152 |
| German Shepherd | 32 | UK Herding | 46 |
| Hungary | 43 |  |  |

#### 2.3.3 Microarray and two-sample comparisons

To assess the effect of variant ascertainment, analysis was repeated based on sites included on the Illumina Canine HD array. Analysis was also performed based on variants that differ between two individuals. For this, a sample from each of the two categories being analyzed was selected from the samples not included in the randomly selected subset used in the sharing analysis. For the breed dogs, comparisons were made between the largest clade, Scenthounds, and the three clades that have the most rare variation.

### 2.4 Human GeoVar analysis

Analysis of human allele sharing was performed to assess the effect of down sampling to 50 individuals per group. Human variation data from the high coverage 1000 Genomes Project sample collection was obtained from [35]. Analysis was limited to passing biallelic variants in the autosomes with no missing data. The same five continental populations used by Biddanda et al. [32] were assessed: AFR (Africa), EUR (Europe), EAS (East Asia), SAS (South Asia), and AMR (admixed Americas). To match the dog sample sizes, 50 individuals were randomly selected from each category. To assess the impact of a variable number of categories, human analysis was also performed with three categories: Africa, Asia (combined EAS and SAS), and European populations.

## 3. Results

### 3.1 Average number of differences between canines

We analyzed patterns of allele sharing among canines based on the genome-wide SNP variation map obtained by the Dog10K consortium [21]. Genetic variants were identified based on alignment of Illumina sequencing reads to the canFam4/UU_Cfam_GSD_1.0 reference assembly derived from a German Shepherd Dog [22]. Following filtering, the analyzed variant data set included 26,585,484 autosomal, biallelic SNPs that were variable among 1,929 samples with a density of ∼120 SNPs/10 kbp (Figure S1), 79% of the original ‘PASS’ SNPs in the data.

The 1000 Genomes Project data reports variation from 2,504 humans [9], with a total of 98,188,417 PASS site SNPs, or about 357 SNPs/10 kb. After performing the same filtering described above, this rate is reduced to about 312 SNPs/10 kb (Figure S2). This represents a 2.6-fold increase over the rate of canine SNPs that were analyzed, with 87% of total PASS SNPs being analyzed.

To assess the genetic differences among samples we determined the number of SNP differences between pairs of samples across categories (Figure 1). As expected, wolves show the greatest amount of genetic variation: the 57 wolf samples differ from each other at a mean of 2,346,972 SNPs. This is followed by village dogs, where the 281 samples show a mean difference of 1,806,946 SNPs. We selected one sample from each of the 321 breeds analyzed, and found a mean difference of 1,702,814 SNPs between breeds. A subset of pairwise breed comparisons show increased SNP distances, around 2 million. This includes comparisons of Japanese breeds with wolfdogs, notably the Czechoslovakian Wolfdog and Saarloos Wolfdog. The breed with the largest mean SNP distance to other breeds is the Shikoku (Table S2). Other breeds that have a high average SNP distance to other breeds include the Norwegian Lundhund, consistent with the extreme population bottleneck this breed experienced [21,36,37].

**Figure 1.**
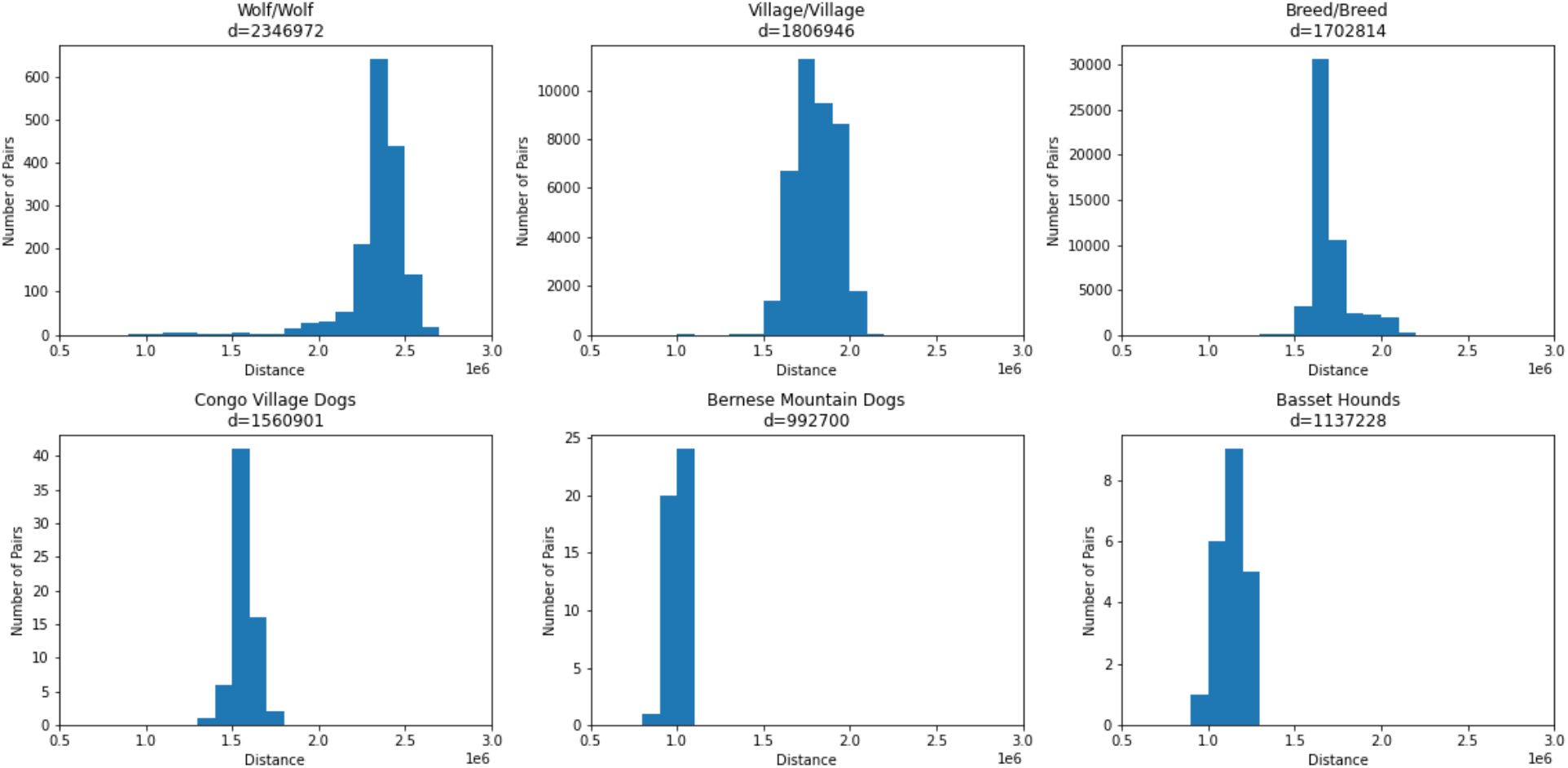
SNP distances between samples. Histograms of SNP distances in groups of wolves, breed dogs, village dogs, village dogs from Congo, Bernese Mountain Dogs and Basset Hounds are shown. The vertical scales are not identical, since there are different counts for different groups. The mean distance between each selection of pairs is given by ‘d’ at the top.

As a comparison, we performed the same analysis comparing samples within a breed. We selected two breeds for comparison: Basset hounds and Bernese Mountain Dogs. We found a mean of 1,137,228 SNP differences among seven Basset hounds and 992,700 SNP differences among ten Bernese Mountain Dogs. To compare this between the difference in a geographic cluster of village dogs, we found that among the 15 Congo Village Dogs, there is a mean SNP difference of 1,560,901 between pairs of samples, consistent with the greater amount of genetic diversity found in village dogs [13,15].

### 3.2 Sharing of alleles among breed dogs, wolves, and village dogs

We constructed GeoVar plots to assess patterns of allele sharing among breed dogs, wolves, and village dogs. These plots offer a visual description of allele sharing as a function of allele frequency in a group and can be considered as a discretized representation of the multi-dimensional site frequency spectrum. Variants are classified as being common (maf ≥ 5%), rare (maf < 5%), or unobserved in each population and plotted based on combinations of categories across sample sets. Since only 50 samples from each category were selected, the minimum possible allele frequency is 1%.

To correct for the uneven size of the groups, we randomly selected 50 breed dogs, wolves, and village dogs (Figure 2). The most common GeoVar classification is for alleles to be common in all three groups: 22% of sites have a minor allele frequency ≥ 5% in all three categories. This is closely followed by sites that are rare in wolves and unobserved in breed dogs and village dogs (21% of sites). An additional ∼10% of sites are common in wolves and unobserved in breed dogs and village dogs, thus ∼31% of sites are only variable in wolves. Approximately 10% of sites are rare in village dogs and unobserved in the other samples, while ∼5% of sites are rare in breed dogs and unobserved elsewhere. To assess the effect of random sample selection, we repeated this process with five different random sample selections (Figure 2). In general, the effect of using a different random sample on the frequency of the categories is small, varying the size of the categories by about +/-1% of the total SNP set, although in certain situations, it does affect the ordering of the categories.

**Figure 2.**
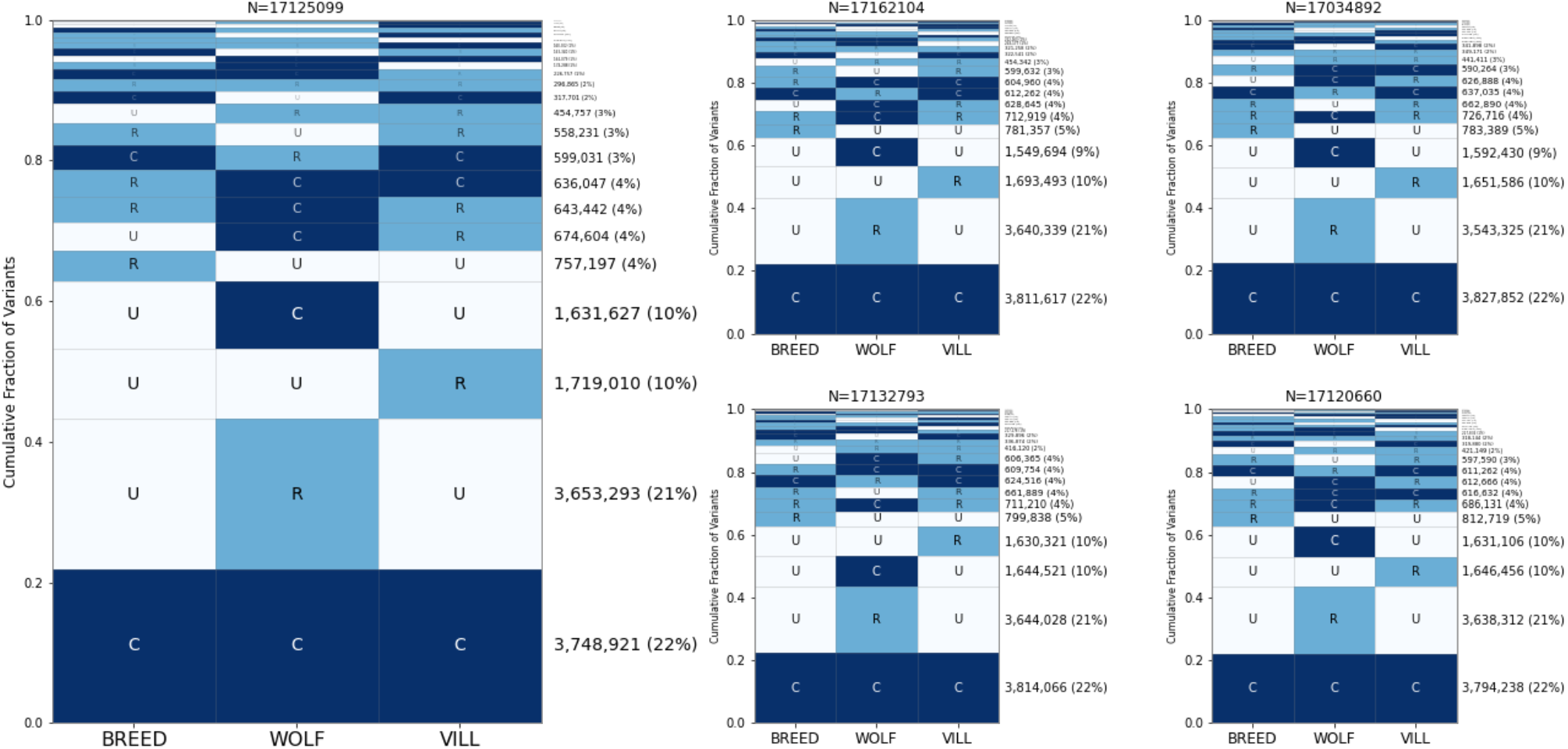
Allele sharing among wolves, village dogs, and breed dogs. The GeoVar plots show the proportion of different patterns of alleles in the three major categories of breed dogs, wolves, and village dogs, shown from left to right. The rows are arranged so that the most common pattern for each single nucleotide variant is at the bottom, with decreasing frequency going toward the top of the figure. Each individual figure represents five separate random samples of 50 canines each, with the larger figure on the left being arbitrarily chosen for easier viewing. The boundary between rare and common is a minor allele frequency < 5%. The total number of SNPs analyzed for each sample set is given at the top of each plot.

Since the effect of analyzing of 50 samples per category versus the hundreds of samples per category used in the human analysis is unclear [32], we repeated the GeoVar analysis of the 1000 Genomes Project samples using a random selection of 50 individuals per group. The analysis in Biddanda *et. al*. found that most variants in humans are rare in a single population and unobserved elsewhere. In our human analyses with a reduced sample size, the most frequent category is variants that are rare in samples from Africa and unobserved elsewhere (Figure S3). Unsurprisingly, with smaller sample sizes, the proportion of variants that are globally common increases, accounting for ∼11% of sites. Since the Dog10K analysis used only three groups instead of the five used for humans, we repeated the analysis by classifying human samples into three groups (Figure S4). This increases the percentage of globally common alleles to ∼15%, still notably lower than the ∼22% found in canines.

### 3.3 Allele sharing within village dogs and breed clades

Next, we assessed allele sharing within sample groups. First, we analyzed village dogs. Based on sample location and Principal Component Analysis we assigned 281 village dog samples into three geographic groups (Figure S5, Table S1): Africa (50 samples from Kenya, Liberia, and Congo), Central Asia (40 samples from Azerbaijan, Bulgaria, Iran, Tajikistan, and Uzbekistan) and East Asia (139 samples from China, Cambodia, Myanmar, and Nepal). We performed GeoVar analysis across these three groups based on a random sampling of 35 individuals per group (Figure 3). The most common pattern was variants that were common in all three groupings, accounting for ∼35% of SNPs. This was followed by variants that were rare in one population but absent in the other three, with the highest proportion of variants present in the village dogs from East Asia.

**Figure 3.**
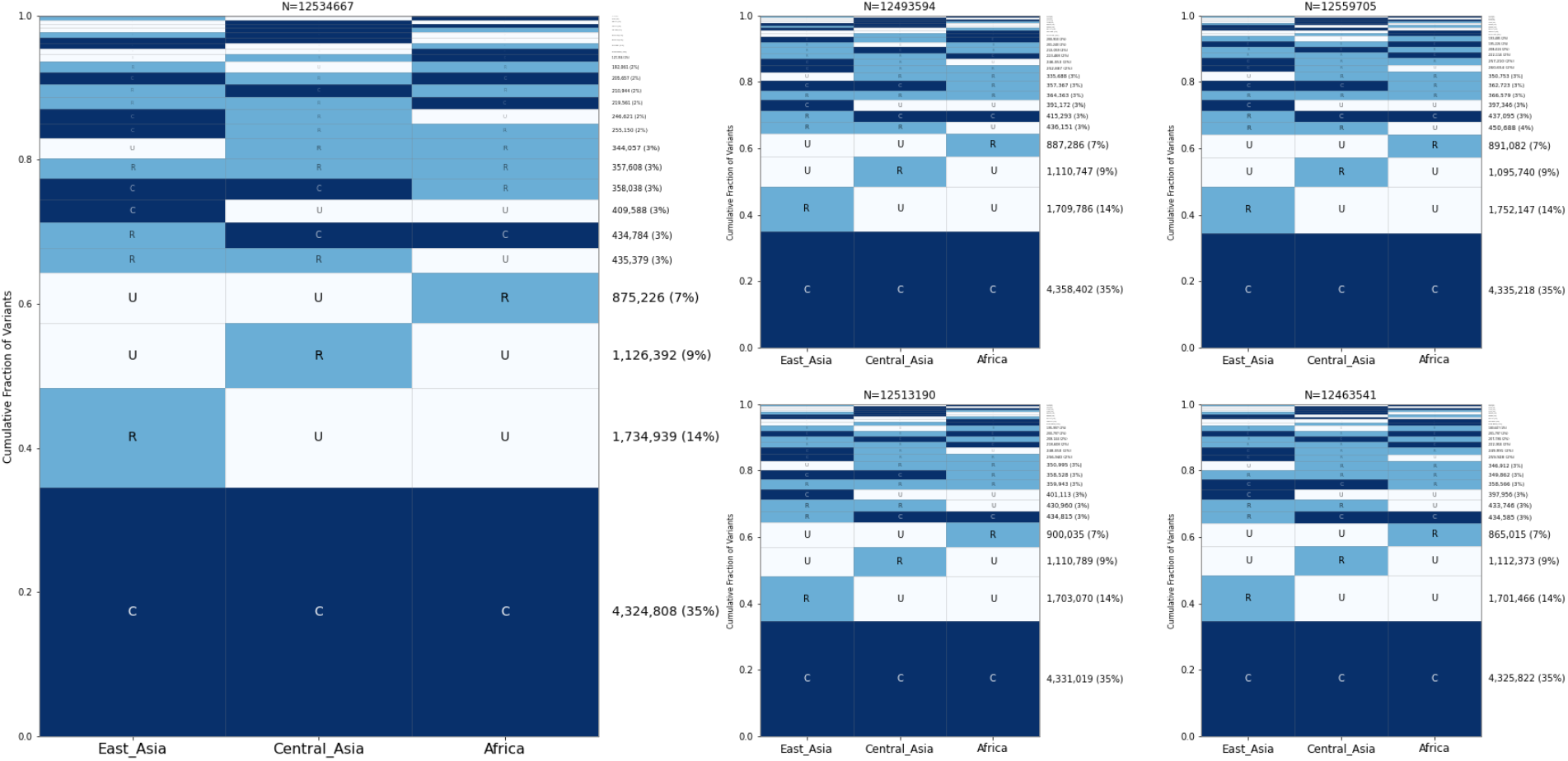
Allele sharing among village dog groups. Five GeoVar plots showing the allele sharing between village dogs based on the geographic groupings, with 35 random samples in each category. Only SNPs which showed polymorphism in village dogs are depicted.

To explore sharing among dog breeds, we analyzed 17 breed clades as defined by the Dog10K project (Table 1) [21]. We selected 30 samples from each clade for analysis and found that variants that are common in all 17 clades were the most common category, accounting for ∼16% of sites (Figure 4). This was followed by variants that were rare in a single clade and absent elsewhere, with the Asian, German Shepherd, and Flockguard Sighthound clades possessing the most rare variation.

**Figure 4.**
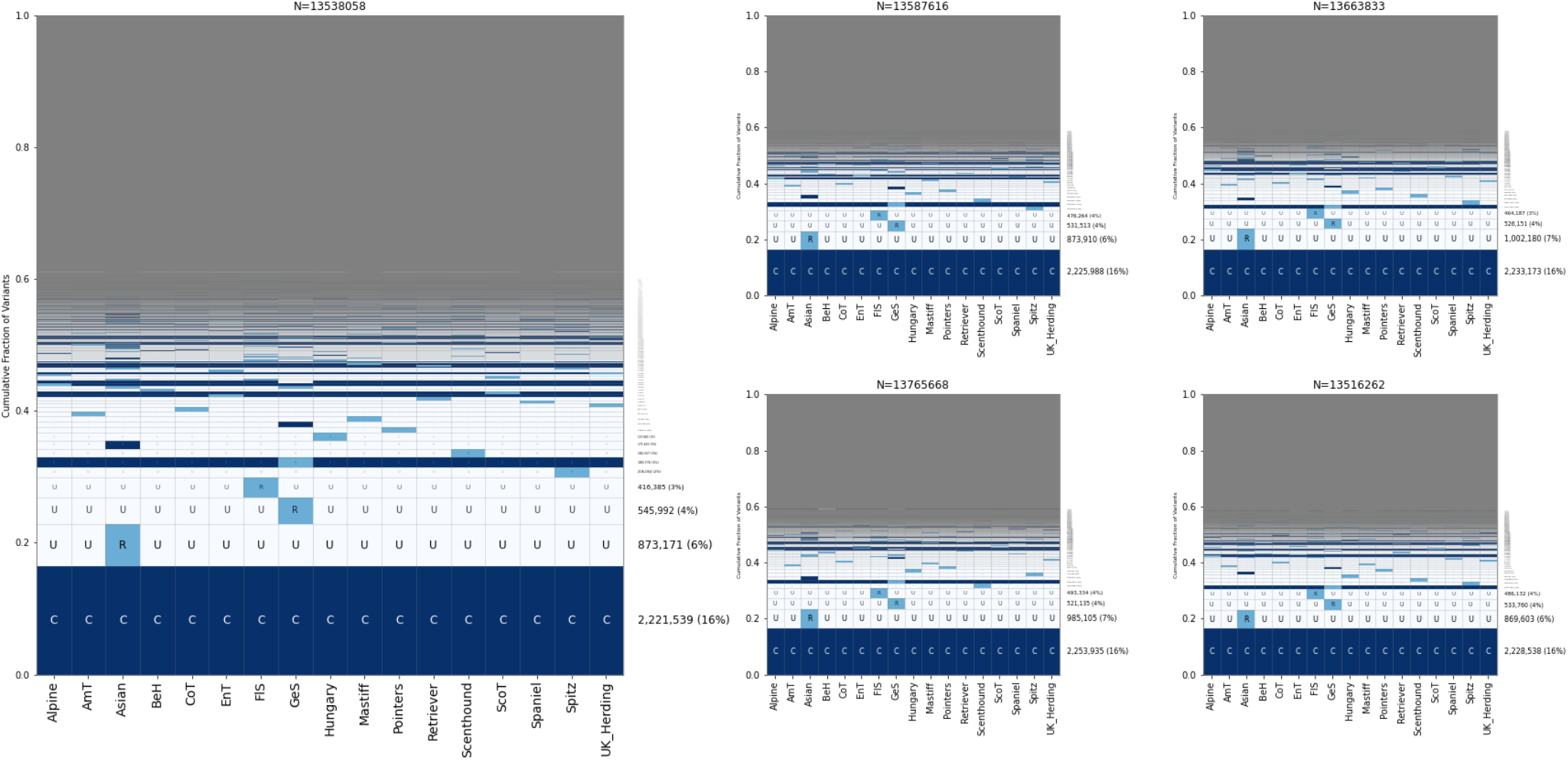
Allele sharing among breed clades. Five GeoVar plots showing allele sharing between breed clades. Only SNPs which show polymorphism in breed dogs are depicted. 30 samples from each population were collected. Abbreviations: AmT=American Terriers, BeH=Belgian Herders, CoT=Contiential Terriers, EnT=English Terriers, FlS=Flockguard Sighthound, GeS=German Shepherd, ScoT=Scottish Terriers. The order of the clades is the same in each plot.

### 3.4 SNP microarrays are skewed toward globally common sites

To assess the effect of ascertainment of sites present on genotyping arrays, we repeated the GeoVar analysis using only the sites present on the Illumina Canine HD SNP array. After filtering, this resulted in 150,299 sites. As expected, the proportion of sites that are common increased dramatically, with ∼65% of all variants being globally common among wolves, village dogs, and wolves (Figure 5). Variants that are rare in one population and common in the others are also more abundant than found in the sequencing data.

**Figure 5.**
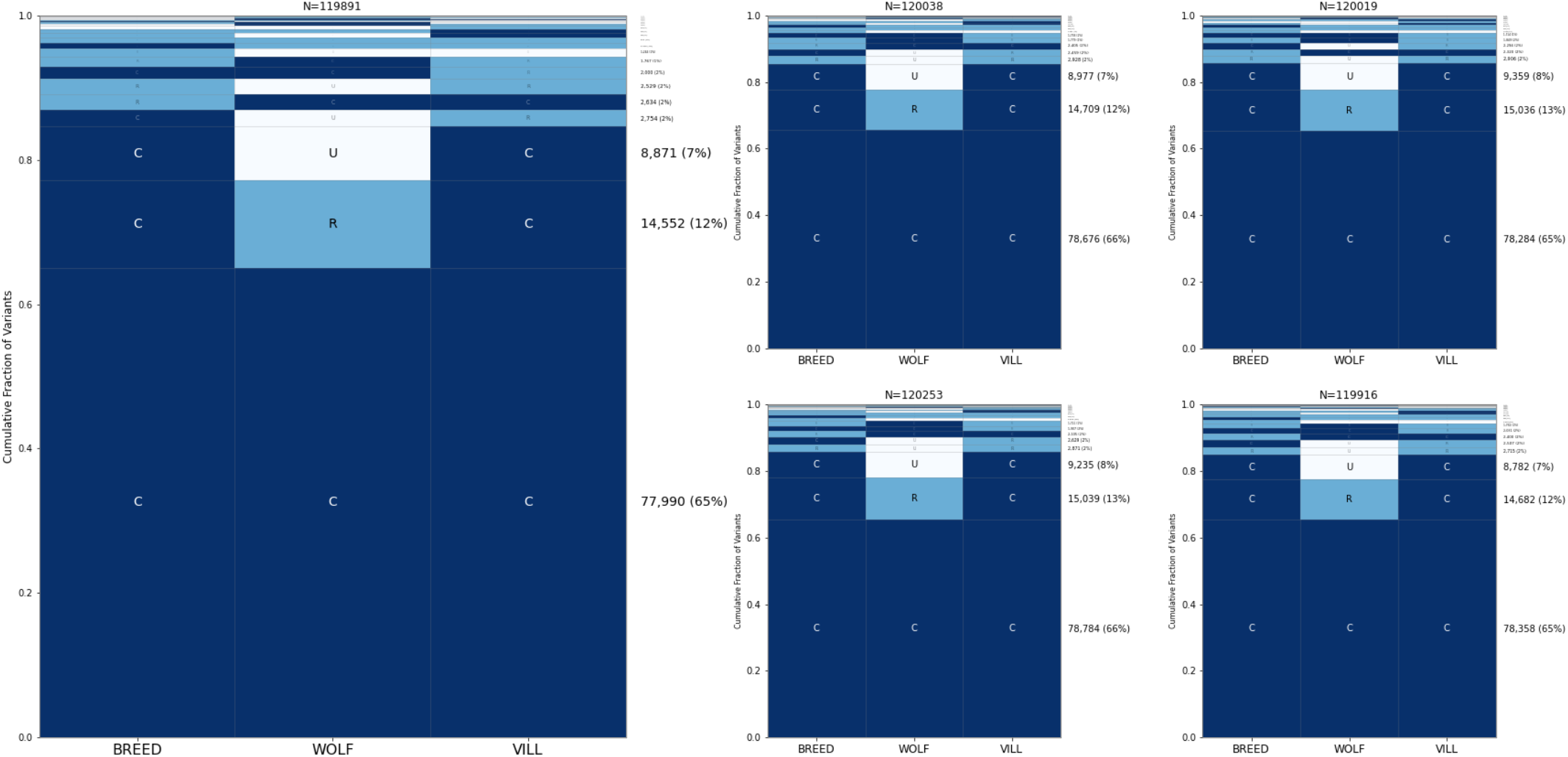
Allele sharing among wolves, village dogs, and breed dogs based on sites present on genotyping arrays. A GeoVar plot using only SNPs found on the Canine HD Illumina array. The five random samples of breeds are the same as in Figure 2.

### 3.5 Common variants account for most of the differences between individual dogs and wolves

To assess the frequency spectrum of sites that differ between two individuals, we repeated GeoVar analysis based on sites that differ between two samples. First, we identified one breed dog (a Golden Retriever), one wolf (from Russia), and one village dog (from Nepal) that were not in the randomly select set used for analysis (which is the same as in the bottom right plot in Figure 3). We then performed GeoVar analysis based on the SNP differences found between each pair of these three samples (Figure 6). Sites that are common in all three groups accounted for ∼56%-73% of the differences between samples.

**Figure 6.**
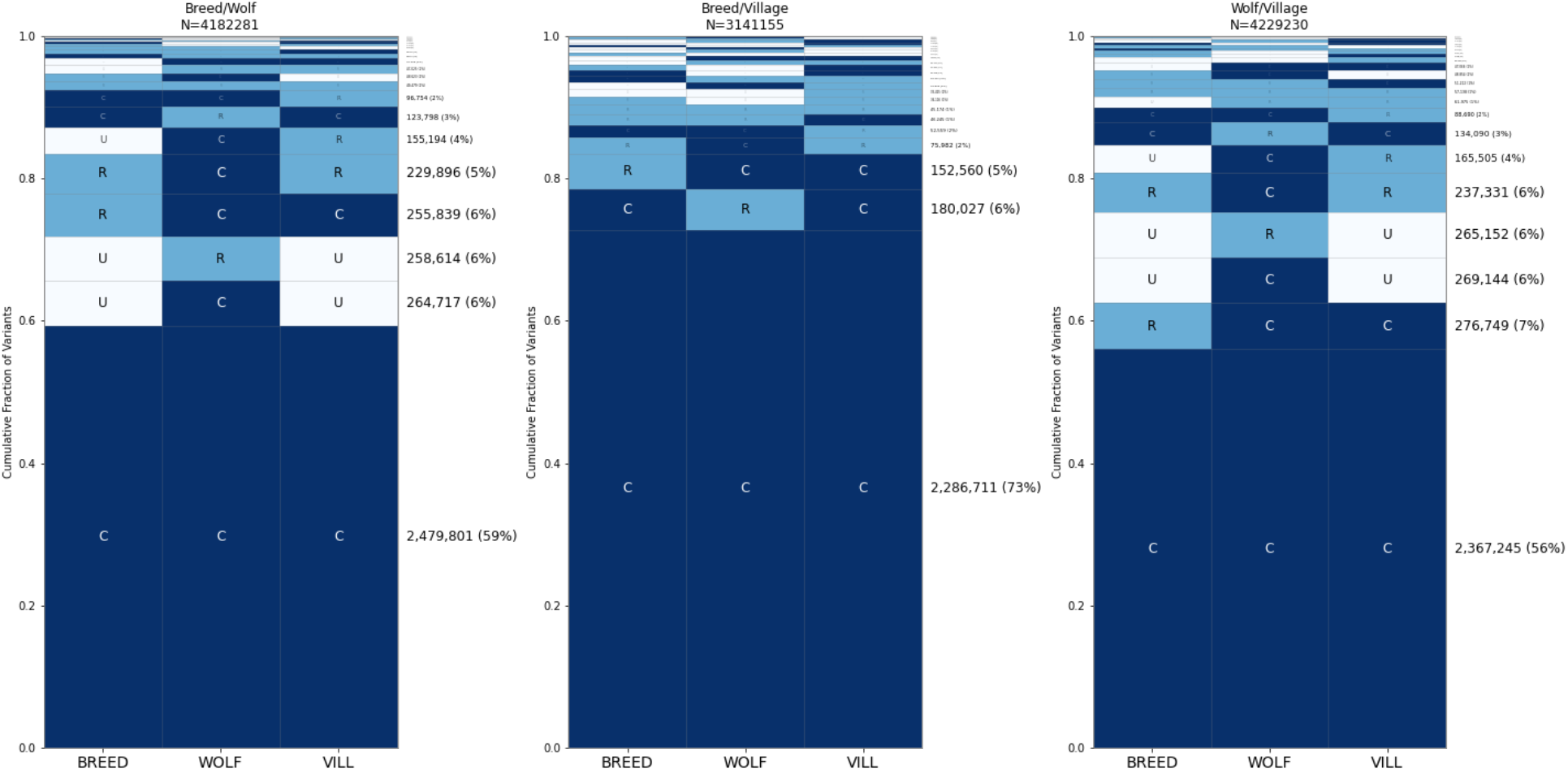
Distribution of alleles that differ between individuals from different categories. A comparison of SNPs that are unique when comparing between breed dogs, wolves, and village dogs is shown. Panel (a) shows SNPs that are different between a breed dog and a wolf, panel (b) shows SNPs that are different between a breed dog and a village dog (Congolese), and panel (c) shows SNPs that are different between a village dog and a wolf. The 50 individuals used in each category are the same as in the bottom right subplot in Figure 4. The two dogs that are used to find differences are not in this random sample. Breed Dog = GOLD000007 (Golden Retriever), Village Dog = VILLNP000001 (Nepal), Wolf=CLUPRU000001 (Russia)

We repeated GeoVar analysis based on sites that differ between two samples belonging to the same group: two breed dogs (a Hellenic Hound and a Swedish White Elkhound), two village dogs (from China and French Polynesia), and two wolves (from Russia and China) (Figure 7). We found that ∼49% of the sites that differ between two wolves are common across the breed dog, village dog, and wolf categories while ∼71% of the sites that differ between two breed dogs or two village dogs are globally common. A similar pattern holds when considering the geographic distribution of variants found between two village dogs, with 71-77% of differences common in each of the East Asia, Central Asia, and Africa groups (Figure 8). Finally, we assessed the distribution of variation across breed clades, finding that ∼49% of the differences between two breed dogs are common across all 17 clades (Figure 9). An additional ∼4% of variants are rare in in the German Shepherd clade but common in all others.

**Figure 7.**
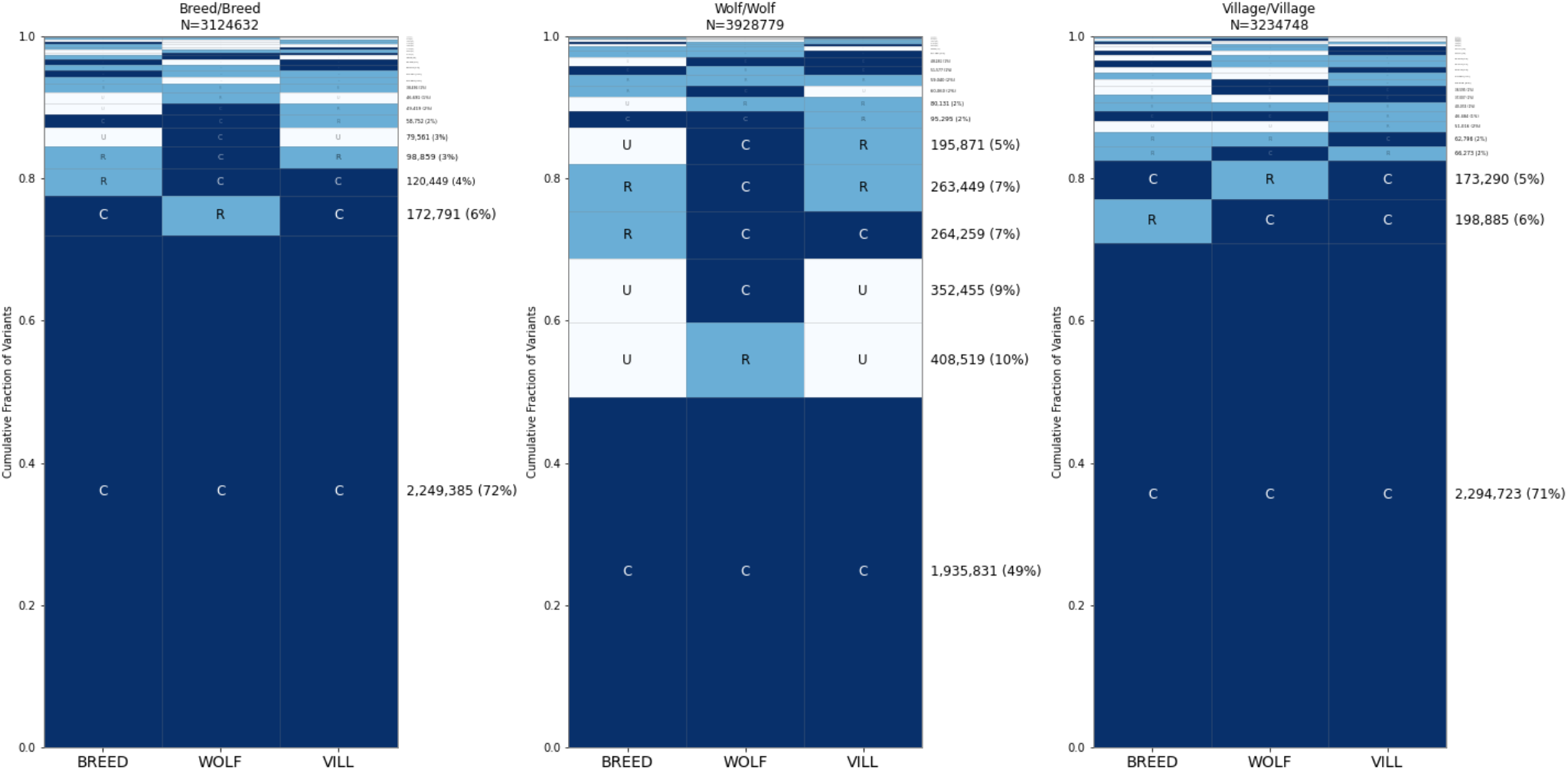
Distribution of alleles that differ between individuals from the same category. The same analysis as in Figure 7, except using two samples from the same category. Breed Dogs = CRTR000009 (Hellenic Hound) and SWWE000006 (Swedish White Elkhound), Wolves = CLUPRU000001 (Russia) and CLUPCN000001 (China), Village Dogs = VILLCN000091 (China), VILLPF000004 (French Polynesia).

**Figure 8.**
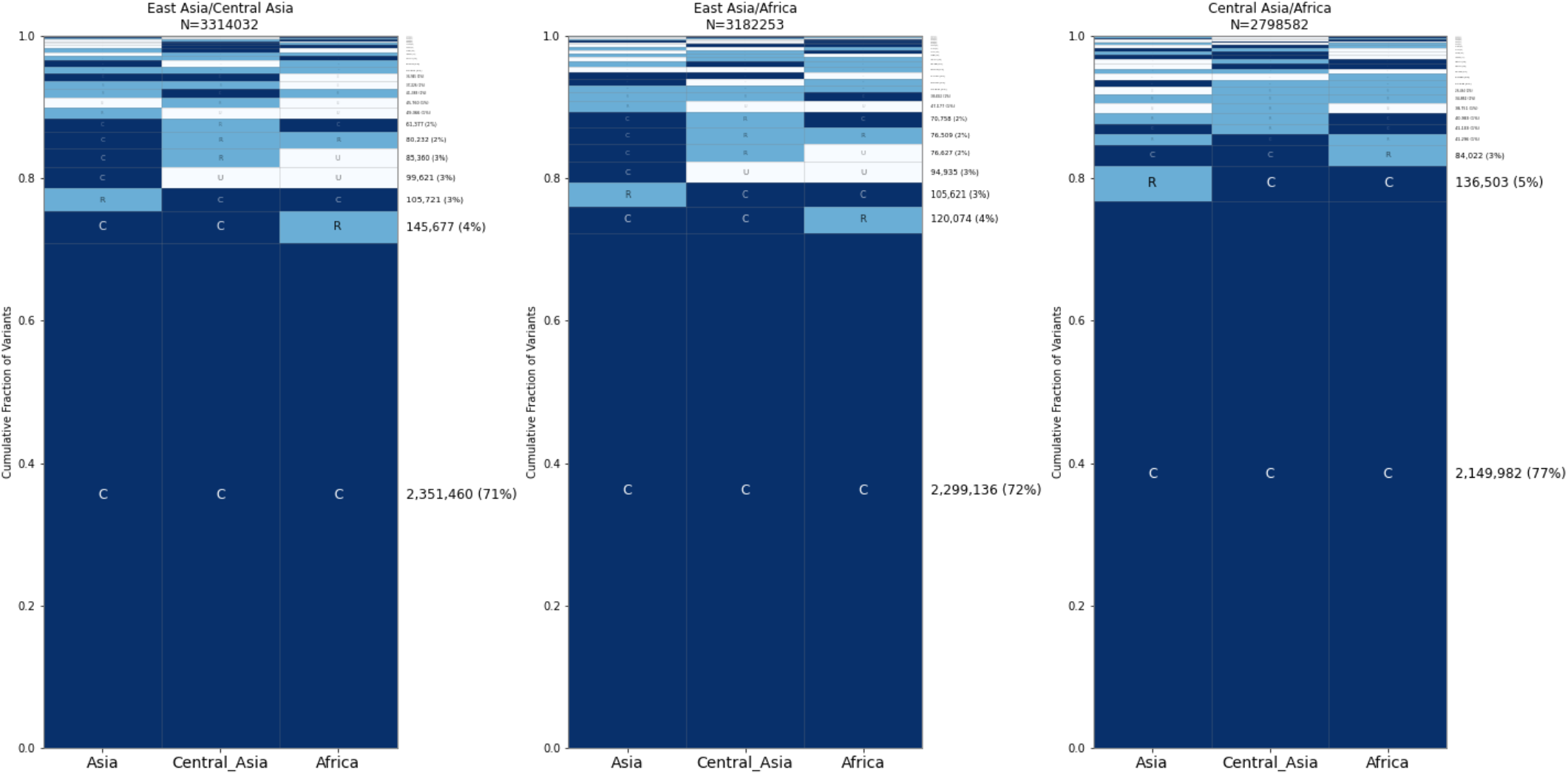
Distribution of alleles that differ between village dogs. Comparisons of SNPs that are different between two village dogs. The dogs used are not found in the random samples of 35 each. Samples used for the comparison: East Asia = VILLCN000099 (China), Central Asia = VILLIR000022 (Iran), Africa = VILLCG000006 (Congo).

**Figure 9.**
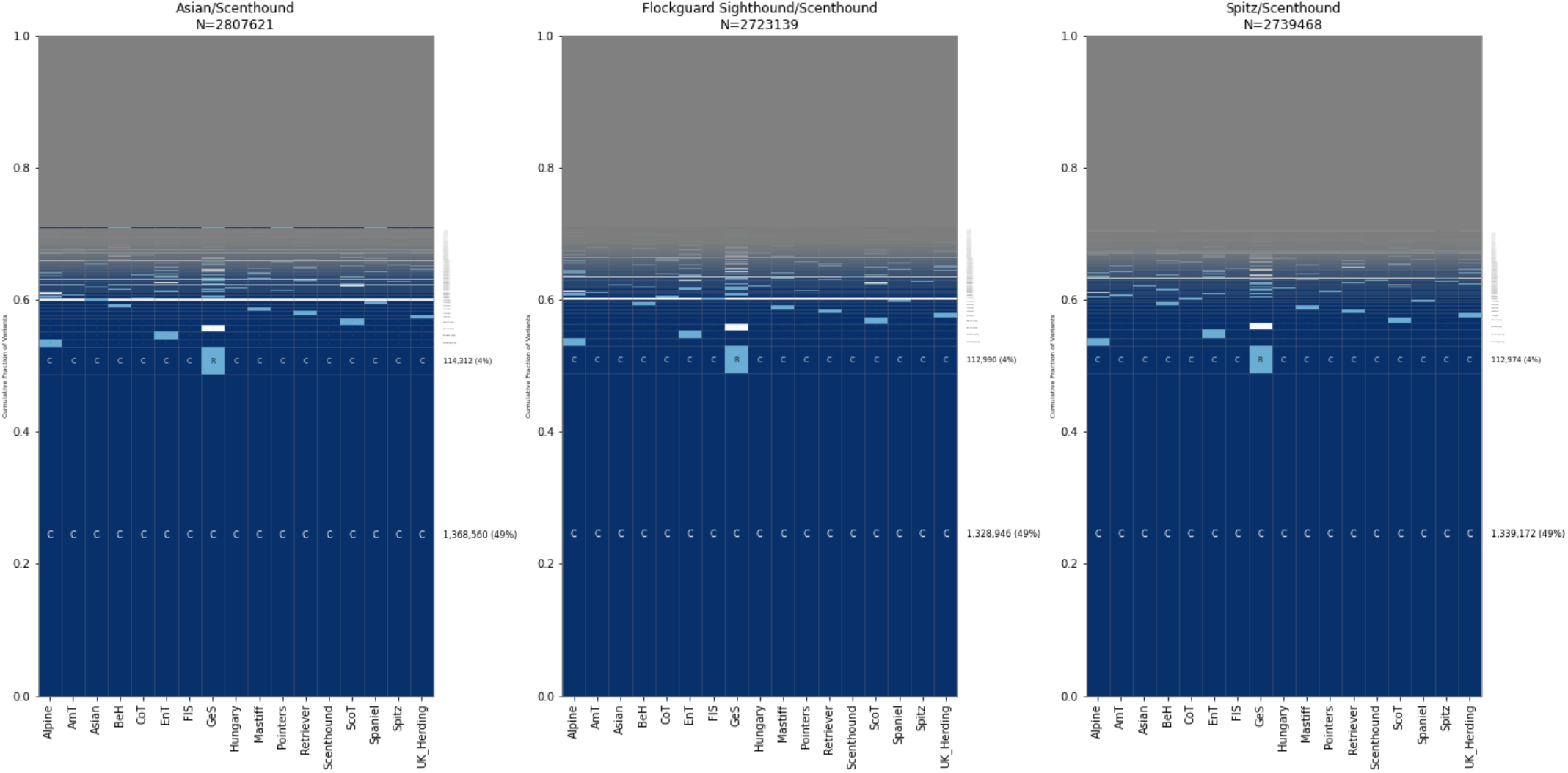
Distribution of alleles that differ between breed dogs. An analysis based on variants that differ between dogs from different breed clades is shown. Comparisons are given for a Scenthound compared to three clades of breed dogs: the Asian clade, Flockguard Sighthound, and Spitz. Abbreviations: AmT=American Terrier, BeH=Belgian Herder, CoT= Continental Terrier, EnT=English Terrier, FlS=Flockguard Sighthound, GeS=German Shepherd, ScoT=Scottish Terrier. Samples used: Scenthound=BMSH0000002, Flockguard Sighthound=CIRN000003, Asian=TIBT000003, Spitz=GSVM000005.

## 4. Discussion

The availability of genome sequencing data from diverse samples allows for an unprecedented view of genetic variation in a species, including variation that is rare or restricted to particular subpopulations. However, the scale of the data sets that are now produced presents barriers for the efficient analysis and visualization of the resulting data, necessitating the use of multiple complementary approaches. In this study, we analyzed patterns of allele sharing among wolves, village dogs, and breed dogs analyzed by the Dog10K consortium [21].

As an initial summary of the data we determined the average SNP distances between samples. As expected, wolves are the most genetically diverse group with an average of ∼2.3 million SNP differences between samples, compared to ∼1.8 million SNPs between village dogs and 1.7 million SNPs between dogs of different breeds. Dogs from the same breed show a lower, but still substantial, amount of diversity, ∼1 million SNPs, although this differs by breeds. We note that these distances are calculated based on SNPs that were identified by Illumina short-read sequencing following a standardized filtering approach [21] and are also reduced by our removal of variants with any missing data across all samples. Direct comparison of high-quality long-read assemblies identifies additional SNP differences [28], as well as other types of variation such as small indels and mobile element insertions that make important contributions to canine genetic diversity [27].

To assess the sharing of SNPs across sample groups we used GeoVar, a visualization method developed for the analysis of human genetic variation [32]. The canine samples we analyzed include fewer individuals per group as well as fewer groupings relative to collections of human genetic variation, resulting in an increased proportion of common alleles. However, when using a sample size of 50 individuals per group, we find that 22% of variants are common across wolves, village dogs, and breed dogs. In comparison ∼15% of alleles are common across three continental groupings of human samples of the same size. This suggests that globally common alleles are approximately 50% more abundant across the three groupings of canines than found in humans.

We found that ∼16% of sites are common across 17 breed clades, followed by variants that are rare in the Asian, German Shepherds, or Flockguard Sighthound clades and unobserved elsewhere. The high proportion of variants that are unique to the Asian clade (∼6%) likely reflects the deep ancestral separation found between dogs across Eurasia [4]. The Dog10K data was generated by aligning reads to the canFam4/UU_Cfam-GSD_1.0 assembly, which was derived from a German Shepherd Dog. Although one may speculate about the effects of reference bias on variation patterns found among German Shepherd-like dogs, we note that similar breed relationship patterns are observed when Illumina sequencing data is analyzed relative to a reference assembly from a Greenland wolf outgroup [38].

We categorized the village dogs into three categories based on PCA and geography. The village dogs showed remarkable similarity between the categories, with ∼35% of alleles being globally common. Village dogs from East Asia tend to have more genetic diversity than village dogs from Central Asia or Africa. However, the differences between the groups are minor.

As expected, most of the differences between any two samples represent variants that are common across all categories. We found that nearly half the sites that differ between two wolves are common across the wolf, village dog, and breed dog categories and that almost half of the differences between two dogs of different breeds are due to sites that are common across all 17 breed clades. Among the village dog categories, nearly three-quarters of the differences are globally common across all three categories.

These results give a more complete picture of how canine variation varies across breeds and between breed dogs, wolves, and village dogs. The GeoVar plots provide a simple visualization that complements other techniques such as SNP distances and PCA. Importantly, GeoVar plots offer a glimpse into the sharing patterns of rare variants that have only been accessible since the advent of large-scale whole genome sequencing studies.

## Supplementary Tables and Figures

Supplementary material for this article is available online, including:

Supplementary Table 1. Village Dog Population Assignments, Supplementary Table 2. Breed Dog SNP Distances, Figure S1. Density of SNPs included in analysis, Figure S2. Density of SNPs from the human 1000 Genomes Project, Figure S3. Human GeoVar analysis with reduced sample sizes, Figure S4. Human GeoVar analysis with three groups, and Figure S5.Principal component analysis of village dogs.

## Author Contributions

M.W. F. and J.M.K conceived of the study, M.W.F. performed analyses, M.W.F. and J.M.K. wrote the manuscript. This research was supported in part through computational resources and services provided by Advanced Research Computing at the University of Michigan, Ann Arbor.

## Data Availability Statement

SNP data from the Dog10K consortium is available in the Zenodo archive at https://zenodo.org/record/8084059. Data from the human 1000 Genomes project high-coverage sequencing is available at https://ftp.1000genomes.ebi.ac.uk/vol1/ftp/data_collections/1000G_2504_high_coverage/working/20190425_NYGC_GATK/.

## Acknowledgements

We thank the members of the Dog10K Consortium for making canine variation data publicly available. We thank Mathew Blacksmith, Emily Koch, Anthony Nguyen, and Peter Schall for helpful feedback on manuscript drafts.

