## Supplementary Tables and Figures for "A variant centric analysis of allele sharing in dogs and wolves"

### Supplementary Table 1. Village Dog Population Assignments

| East Asia | | | | Central Asia | Africa |
|---|---|---|---|---|---|
| VILLCN000001 | VILLCN000002 | VILLCN000003 | VILLCN000004 | VILLAZ000001 | VILLCG000001 |
| VILLCN000005 | VILLCN000006 | VILLCN000007 | VILLCN000008 | VILLAZ000002 | VILLCG000002 |
| VILLCN000009 | VILLCN000011 | VILLCN000012 | VILLCN000013 | VILLAZ000003 | VILLCG000003 |
| VILLCN000014 | VILLCN000015 | VILCN000016 | VILLCN000018 | VILLAZ000004 | VILLCG000004 |
| VILLCN000019 | VILLCN000020 | VILLCN000021 | VILLCN000023 | VILLAZ000005 | VILLCG000005 |
| VILLCN000025 | VILLCN000026 | VILLCN000027 | VILLCN000028 | VILLAZ000006 | VILLCG000006 |
| VILLCN000030 | VILLCN000031 | VILCN000032 | VILLCN000034 | VILLAZ000007 | VILLCG000007 |
| VILLCN000035 | VILLCN000036 | VILLCN000037 | VILLCN000038 | VILLIR000001 | VILLCG000008 |
| VILLCN000039 | VILLCN000040 | VILLCN000042 | VILLCN000044 | VILLIR000002 | VILLCG000009 |
| VILLCN000046 | VILLCN000047 | VILLCN000048 | VILLCN000049 | VILLIR000003 | VILLCG000010 |
| VILLCN000050 | VILLCN000051 | VILLCN000052 | VILLCN000053 | VILLIR000004 | VILLCG000011 |
| VILLCN000054 | VILLCN000055 | VILLCN000056 | VILLCN000057 | VILLIR000005 | VILLCG000012 |
| VILLCN000058 | VILLCN000059 | VILLCN000060 | VILLCN000062 | VILLIR000006 | VILLCG000013 |
| VILLCN000063 | VILLCN000064 | VILLCN000065 | VILLCN000066 | VILLIR000007 | VILLCG000014 |
| VILLCN000068 | VILLCN000069 | VILLCN000070 | VILLCN000071 | VILLIR000008 | VILLCG000015 |
| VILLCN000072 | VILLCN000073 | VILLCN000074 | VILLCN000075 | VILLIR000009 | VILLCG000016 |
| VILLCN000078 | VILLCN000079 | VILLCN000080 | VILLCN000081 | VILLIR000011 | VILLKE000001 |
| VILLCN000084 | VILLCN000086 | VILLCN000087 | VILLCN000088 | VILLIR000012 | VILLKE000002 |
| VILLCN000089 | VILLCN000090 | VILLCN000091 | VILLCN000092 | VILLIR000013 | VILLKE000003 |
| VILLCN000093 | VILLCN000094 | VILLCN000097 | VILLCN000099 | VILLIR000014 | VILLKE000004 |
| VILLCN000100 | VILLCN000103 | VILLCN000104 | VILLCN000108 | VILLIR000015 | VILLKE000005 |
| VILLCN000109 | VILLCN000111 | VILLCN000112 | VILLCN000114 | VILLIR000016 | VILLKE000006 |
| VILLCN000115 | VILLCN000116 | VILLCN000117 | VILLCN000118 | VILLIR000017 | VILLKE000007 |
| VILLCN000120 | VILLCN000121 | VILLCN000123 | VILLCN000124 | VILLIR000018 | VILLKE000008 |
| VILLCN000125 | VILLCN000126 | VILLCN000128 | VILLCN000129 | VILLIR000019 | VILLKE000009 |
| VILLCN000130 | VILLCN000131 | VILLCN000132 | VILLCN000133 | VILLIR000020 | VILLKE000010 |
| VILLCN000134 | VILLCN000135 | VILLCN000136 | VILLCN000137 | VILLIR000021 | VILLKE000011 |
| VILLCN000139 | VILLCN000140 | VILLCN000142 | VILLCN000143 | VILLIR000022 | VILLKE000012 |
| VILLCN000144 | VILLCN000145 | VILLCN000146 | VILLCN000147 | VILLIR000023 | VILLKE000013 |
| VILLCN000148 | VILLCN000152 | VILLCN000153 | VILLCN000154 | VILLIR000025 | VILLKE000014 |
| VILLKH000001 | VILLKH000002 | VILLKH000003 | VILLKH000004 | VILLIR000026 | VILLKE000015 |
| VILLMM000004 | VILLMM000005 | VILLNP000001 | VILLNP000002 | VILLIR000028 | VILLKE000016 |
| VILLNP000003 | VILLNP000004 | VILLNP000005 | VILLNP000006 | VILLIR000030 | VILLKE000017 |
| VILLNP000007 | VILLNP000008 | VILLTH00002 | VILLTH000005 | VILLTJ000001 | VILLKE000019 |
| VILLTH000006 | VILLTH000008 | VILLTH000009 | | VILLUZ000001 | VILLLR000001 |
| | | | | VILLUZ000002 | VILLLR000002 |
| | | | | VILLUZ000003 | VILLLR000003 |
| | | | | VILLUZ000004 | VILLLR000004 |
| | | | | VILLUZ000005 | VILLLR000005 |
| | | | | VILLUZ000006 | VILLLR000006 |
| | | | | | VILLLR000007 |
| | | | | | VILLIR000008 |
| | | | | | VILLLR000009 |
| | | | | | VILLLR000010 |
| | | | | | VILLLR000011 |
| | | | | | VILLLR000012 |
| | | | | | VILLLR000013 |
| | | | | | VILLLR000014 |
| | | | | | VILLLR000015 |
| | | | | | VILLLR000016 |

**Supplementary Table 2. Breed Dog SNP Distances**

SNP distances were calculated between one dog randomly selected from each analyzed breed. The minimum, maximum, mean, and median results of the comparisons are shown for each breed.

| Breed Name | Minimum | Mean | Median | Maximum |
|---|---|---|---|---|
| Cocker Spaniel | 1,561,755 | 1,694,320 | 1,671,977 | 2,115,342 |
| American Eskimo Dog | 1,590,627 | 1,686,893 | 1,668,605 | 2,092,240 |
| Affenpinscher | 1,533,830 | 1,697,714 | 1,675,342 | 2,105,508 |
| Afghan Hound | 1,723,167 | 1,837,167 | 1,825,825 | 2,141,371 |
| American Foxhound | 1,437,571 | 1,652,102 | 1,637,385 | 2,091,209 |
| American Hairless Terrier | 1,570,635 | 1,675,535 | 1,653,567 | 2,083,619 |
| Airedale Terrier | 1,662,203 | 1,746,878 | 1,723,741 | 2,159,507 |
| Akbash | 1,626,080 | 1,728,518 | 1,713,258 | 2,080,192 |
| Japanese Akita | 1,438,805 | 2,024,604 | 2,026,653 | 2,227,931 |
| American Akita | 1,438,805 | 1,997,714 | 1,999,429 | 2,203,001 |
| Alpine Dachsbracke | 1,594,575 | 1,686,243 | 1,664,297 | 2,098,755 |
| Alaskan Malamute | 1,512,856 | 1,918,786 | 1,915,894 | 2,138,102 |
| American Bulldog | 1,391,027 | 1,664,364 | 1,648,888 | 2,096,842 |
| American Staffordshire Terrier | 1,464,659 | 1,656,107 | 1,639,106 | 2,098,154 |
| American Water Spaniel | 1,535,998 | 1,674,361 | 1,653,804 | 2,099,800 |
| Anatolian Shepherd Dog | 1,644,556 | 1,766,674 | 1,752,345 | 2,102,797 |
| Appenzeller Sennenhund | 1,554,004 | 1,748,469 | 1,726,667 | 2,145,059 |
| Artois Hound | 1,515,284 | 1,690,477 | 1,673,187 | 2,119,880 |
| Ariegeois | 1,430,344 | 1,626,185 | 1,606,564 | 2,073,452 |
| Ariege Pointer | 1,483,614 | 1,669,549 | 1,650,505 | 2,102,592 |
| Silky Terrier | 1,536,733 | 1,681,655 | 1,659,247 | 2,101,459 |
| Australian Cattle Dog | 1,606,202 | 1,697,732 | 1,676,907 | 2,101,109 |
| Auvergne Pointer | 1,535,668 | 1,654,623 | 1,633,111 | 2,090,715 |
| Australian Kelpie | 1,575,297 | 1,669,692 | 1,647,102 | 2,092,434 |
| Australian Shepherd | 1,575,895 | 1,680,296 | 1,657,620 | 2,088,079 |
| Australian Terrier | 1,538,012 | 1,685,469 | 1,664,701 | 2,091,833 |
| Basset Artesien Normand | 1,438,154 | 1,697,190 | 1,677,850 | 2,118,607 |
| Basset Hound | 1,438,154 | 1,683,399 | 1,664,963 | 2,101,859 |
| White Swiss Shepherd Dog | 1,162,708 | 1,682,604 | 1,672,702 | 2,101,087 |
| Beauceron | 1,581,119 | 1,670,400 | 1,648,748 | 2,106,902 |
| Belgian Tervuren | 1,378,416 | 1,664,012 | 1,644,161 | 2,081,946 |
| Bedlington Terrier | 1,630,948 | 1,723,028 | 1,699,071 | 2,136,263 |
| Belgian Sheepdog | 1,100,241 | 1,694,446 | 1,674,037 | 2,107,270 |
| Bergamasco Sheepdog | 1,560,095 | 1,651,025 | 1,630,440 | 2,082,451 |
| Basset Fauve de Bretagne | 1,466,045 | 1,662,397 | 1,642,150 | 2,092,049 |
| Blue Gascony Griffon | 1,430,344 | 1,621,259 | 1,604,908 | 2,065,223 |
| Bohemian Shepherd | 1,189,208 | 1,729,101 | 1,719,125 | 2,132,483 |
| Bichon Frise | 1,580,008 | 1,688,462 | 1,667,066 | 2,103,585 |
| Havanese | 1,495,989 | 1,656,045 | 1,636,221 | 2,075,068 |
| Billy | 1,195,380 | 1,659,373 | 1,646,568 | 2,085,884 |
| Bloodhound | 1,509,127 | 1,691,883 | 1,674,750 | 2,118,099 |
| Russian Tsvetnaya Bolonka | 1,613,633 | 1,695,358 | 1,677,869 | 2,073,532 |
| Blue Picardy Spaniel | 1,279,443 | 1,659,718 | 1,640,192 | 2,090,311 |

| | | | | |
|---|---|---|---|---|
| **Bluetick Coonhound** | 1,480,782 | 1,648,809 | 1,628,257 | 2,085,776 |
| **Belgian Malinois** | 1,418,225 | 1,655,842 | 1,637,280 | 2,086,034 |
| **Bavarian Mountain Scent Hound** | 1,542,663 | 1,672,418 | 1,650,953 | 2,096,019 |
| **Bouvier des Ardennes** | 1,508,874 | 1,657,091 | 1,637,853 | 2,075,391 |
| **Boerboel** | 1,417,226 | 1,646,636 | 1,626,805 | 2,086,394 |
| **Bolognese** | 1,545,256 | 1,657,826 | 1,638,938 | 2,080,650 |
| **Bourbonnais Pointing Dog** | 1,502,749 | 1,673,108 | 1,653,634 | 2,113,155 |
| **Boston Terrier** | 1,473,671 | 1,659,711 | 1,641,234 | 2,098,397 |
| **Bouvier des Flandres** | 1,582,781 | 1,689,535 | 1,665,215 | 2,109,776 |
| **Boxer** | 1,406,464 | 1,719,292 | 1,704,845 | 2,145,332 |
| **Boykin Spaniel** | 1,535,998 | 1,674,030 | 1,651,449 | 2,099,669 |
| **Bracco Italiano** | 1,483,614 | 1,685,613 | 1,667,189 | 2,104,935 |
| **Barbet** | 1,561,067 | 1,655,433 | 1,632,890 | 2,090,879 |
| **Braques Francais** | 1,443,750 | 1,640,969 | 1,620,523 | 2,082,922 |
| **Briquet Griffon Vendeen** | 1,318,368 | 1,652,911 | 1,636,843 | 2,089,797 |
| **Broholmer** | 1,554,128 | 1,693,555 | 1,670,083 | 2,114,599 |
| **Briard** | 1,581,119 | 1,688,851 | 1,666,237 | 2,112,870 |
| **Brittany** | 1,491,890 | 1,661,835 | 1,642,919 | 2,087,104 |
| **Bruno Jura Hound** | 1,458,197 | 1,653,931 | 1,633,249 | 2,092,756 |
| **Bernese Mountain Dog** | 1,547,768 | 1,693,370 | 1,668,947 | 2,101,308 |
| **Black Russian Terrier** | 1,522,173 | 1,668,695 | 1,647,280 | 2,086,298 |
| **Brussels Griffon** | 1,115,238 | 1,685,894 | 1,664,499 | 2,099,124 |
| **Brazilian Terrier** | 1,546,312 | 1,667,549 | 1,645,103 | 2,099,597 |
| **Black and Tan Coonhound** | 1,536,950 | 1,683,232 | 1,663,581 | 2,104,593 |
| **French Bulldog** | 1,453,522 | 1,675,719 | 1,658,800 | 2,109,543 |
| **English Bulldog** | 1,037,054 | 1,690,575 | 1,679,717 | 2,121,233 |
| **Bullmastiff** | 1,404,997 | 1,696,800 | 1,678,507 | 2,122,647 |
| **Bull Terrier** | 785,014 | 1,732,413 | 1,718,354 | 2,154,921 |
| **Biewer Terrier** | 1,370,539 | 1,678,674 | 1,655,855 | 2,106,140 |
| **Canaan Dog** | 1,672,740 | 1,765,371 | 1,750,843 | 2,111,670 |
| **Cairn Terrier** | 1,588,516 | 1,683,524 | 1,661,465 | 2,103,382 |
| **Dogo Canario** | 1,455,013 | 1,649,083 | 1,630,599 | 2,092,889 |
| **Cardigan Welsh Corgi** | 1,525,609 | 1,667,733 | 1,646,342 | 2,090,284 |
| **Schipperke** | 1,625,671 | 1,706,206 | 1,685,230 | 2,100,429 |
| **Central Asian Shepherd Dog** | 1,689,965 | 1,778,310 | 1,766,056 | 2,080,962 |
| **Caucasian Ovcharka** | 1,626,508 | 1,721,673 | 1,706,864 | 2,079,431 |
| **Curly-Coated Retriever** | 1,603,604 | 1,693,264 | 1,669,355 | 2,105,630 |
| **Cesky Fousek** | 1,423,240 | 1,643,719 | 1,623,610 | 2,088,007 |
| **Chesapeake Bay Retriever** | 1,545,676 | 1,645,368 | 1,623,026 | 2,073,199 |
| **Chihuahua** | 1,570,526 | 1,668,122 | 1,649,101 | 2,076,583 |
| **Japanese Chin** | 1,742,654 | 1,811,800 | 1,800,000 | 2,071,005 |
| **Chow Chow** | 1,803,537 | 2,051,457 | 2,054,853 | 2,241,064 |
| **Cirneco dell'Etna** | 1,511,162 | 1,687,299 | 1,666,587 | 2,101,063 |
| **Cavalier King Charles Spaniel** | 1,241,321 | 1,710,505 | 1,689,468 | 2,131,413 |
| **Clumber Spaniel** | 1,626,628 | 1,732,527 | 1,709,892 | 2,149,045 |
| **Cane Corso** | 1,434,395 | 1,653,388 | 1,635,616 | 2,081,034 |
| **Continental Bulldog** | 1,037,054 | 1,668,280 | 1,657,551 | 2,105,760 |

| | | | | |
|---|---|---|---|---|
| **Collie** | 1,305,666 | 1,731,757 | 1,712,324 | 2,135,479 |
| **Chinook** | 1,524,614 | 1,765,515 | 1,751,152 | 2,117,645 |
| **Coton de Tulear** | 1,586,772 | 1,672,677 | 1,651,347 | 2,075,975 |
| **Chinese Crested** | 1,582,238 | 1,674,129 | 1,654,009 | 2,085,003 |
| **Croatian Sheepdog** | 1,569,870 | 1,659,602 | 1,639,508 | 2,075,685 |
| **Cretan Tracer** | 1,574,775 | 1,675,836 | 1,657,390 | 2,082,306 |
| **Hellenic Hound** | 1,563,836 | 1,664,244 | 1,646,449 | 2,073,639 |
| **Catalan Sheepdog** | 1,592,756 | 1,675,202 | 1,653,712 | 2,088,476 |
| **Cesky Terrier** | 1,532,756 | 1,736,933 | 1,714,774 | 2,141,374 |
| **Catalburun** | 1,534,037 | 1,640,893 | 1,623,079 | 2,070,533 |
| **Czechoslovakian Wolfdog** | 1,565,861 | 1,962,599 | 1,960,198 | 2,286,807 |
| **Dachshund** | 1,573,066 | 1,663,161 | 1,641,739 | 2,086,515 |
| **Dalmatian** | 1,582,955 | 1,675,805 | 1,653,283 | 2,098,612 |
| **Dandie Dinmont Terrier** | 1,630,948 | 1,710,712 | 1,688,315 | 2,121,039 |
| **Dogo Argentino** | 1,539,827 | 1,685,106 | 1,665,854 | 2,093,115 |
| **Dogue de Bordeaux** | 1,519,635 | 1,693,230 | 1,675,919 | 2,108,798 |
| **Scottish Deerhound** | 1,588,522 | 1,717,737 | 1,694,975 | 2,136,517 |
| **Doberman Pinscher** | 1,644,450 | 1,722,249 | 1,701,225 | 2,121,027 |
| **Drever** | 1,589,861 | 1,685,101 | 1,663,113 | 2,095,039 |
| **Danish-Swedish Farmdog** | 1,514,772 | 1,636,866 | 1,614,851 | 2,075,159 |
| **Dutch Shepherd** | 1,505,805 | 1,681,959 | 1,662,086 | 2,093,386 |
| **Dutch Partridge Dog** | 1,587,443 | 1,697,549 | 1,675,434 | 2,114,743 |
| **English Cocker Spaniel** | 1,465,573 | 1,688,588 | 1,665,828 | 2,117,624 |
| **East-European shepherd** | 1,113,194 | 1,670,329 | 1,662,429 | 2,092,763 |
| **English Foxhound** | 1,291,903 | 1,676,730 | 1,665,398 | 2,111,897 |
| **English Toy Spaniel** | 1,241,321 | 1,692,779 | 1,671,288 | 2,109,739 |
| **English Shepherd** | 1,553,894 | 1,655,736 | 1,634,752 | 2,075,702 |
| **English Toy Terrier** | 1,444,568 | 1,670,315 | 1,647,968 | 2,103,438 |
| **Elo** | 1,587,233 | 1,744,434 | 1,734,779 | 2,004,826 |
| **Entlebucher Mountain Dog** | 1,554,004 | 1,741,456 | 1,719,467 | 2,140,338 |
| **English Pointer** | 1,510,478 | 1,690,356 | 1,671,281 | 2,125,552 |
| **English Setter** | 1,479,729 | 1,687,559 | 1,667,456 | 2,118,395 |
| **Estrela Mountain Dog** | 1,568,368 | 1,660,610 | 1,640,543 | 2,075,570 |
| **English Springer Spaniel** | 1,590,399 | 1,686,545 | 1,663,413 | 2,105,343 |
| **Estonian Hound** | 1,547,273 | 1,670,729 | 1,650,581 | 2,089,888 |
| **Eurasier** | 1,587,233 | 1,845,376 | 1,838,829 | 2,055,724 |
| **Field Spaniel** | 1,465,573 | 1,702,796 | 1,680,451 | 2,128,243 |
| **Finnish Hound** | 1,553,232 | 1,678,386 | 1,659,350 | 2,087,701 |
| **Finnish Spitz** | 1,622,738 | 1,758,236 | 1,743,467 | 2,088,917 |
| **Flat-Coated Retriever** | 1,566,636 | 1,710,036 | 1,686,133 | 2,123,911 |
| **Formosan Mountain Dog** | 1,781,560 | 1,856,148 | 1,849,386 | 2,063,321 |
| **French Spaniel** | 1,567,236 | 1,676,550 | 1,653,750 | 2,108,038 |
| **Wire Fox Terrier** | 1,269,032 | 1,708,310 | 1,690,091 | 2,121,978 |
| **Smooth Fox Terrier** | 1,452,735 | 1,703,677 | 1,684,018 | 2,119,560 |
| **Great Anglo-French Tricolour Hound** | 1,229,958 | 1,673,511 | 1,660,171 | 2,097,550 |
| **Galgo Espanol** | 1,522,177 | 1,649,698 | 1,629,102 | 2,073,594 |
| **Great Anglo-French White and Orange Hound** | 1,195,380 | 1,643,994 | 1,633,441 | 2,087,306 |

| | | | | |
|---|---|---|---|---|
| **Grand Bleu de Gascogne** | 1,413,871 | 1,654,487 | 1,636,958 | 2,096,851 |
| **Grand Basset Griffon Vendeen** | 1,318,161 | 1,681,808 | 1,661,637 | 2,108,880 |
| **Griffon Fauve de Bretagne** | 1,466,045 | 1,636,848 | 1,616,745 | 2,075,233 |
| **Grand Griffon Vendeen** | 1,318,368 | 1,665,845 | 1,651,719 | 2,098,685 |
| **German Hound** | 1,603,460 | 1,701,587 | 1,678,520 | 2,116,864 |
| **German Hunting Terrier** | 1,465,455 | 1,661,826 | 1,642,639 | 2,092,039 |
| **Glen of Imaal Terrier** | 1,543,988 | 1,687,440 | 1,666,739 | 2,111,919 |
| **German Spaniel** | 1,601,168 | 1,689,753 | 1,668,080 | 2,104,459 |
| **Golden Retriever** | 1,575,976 | 1,678,466 | 1,655,233 | 2,097,228 |
| **Gordon Setter** | 1,504,571 | 1,653,029 | 1,632,959 | 2,090,477 |
| **German Pinscher** | 1,502,220 | 1,680,592 | 1,658,244 | 2,102,027 |
| **Great Pyrenees** | 1,527,484 | 1,706,198 | 1,686,358 | 2,111,346 |
| **Greenland Dog** | 1,512,856 | 1,959,934 | 1,958,030 | 2,160,796 |
| **Groenendael** | 1,100,241 | 1,703,658 | 1,684,623 | 2,110,020 |
| **Griffon Nivernais** | 1,469,782 | 1,635,673 | 1,618,415 | 2,070,582 |
| **German Shepherd Dog** | 1,113,194 | 1,674,329 | 1,666,857 | 2,097,873 |
| **German Shorthaired Pointer** | 1,443,750 | 1,650,253 | 1,631,075 | 2,098,229 |
| **Greater Swiss Mountain Dog** | 1,547,768 | 1,728,133 | 1,707,022 | 2,138,509 |
| **Giant Schnauzer** | 1,522,173 | 1,669,596 | 1,647,075 | 2,086,089 |
| **German Spitz Klein** | 1,423,291 | 1,659,550 | 1,640,558 | 2,065,213 |
| **German Spitz** | 1,413,425 | 1,678,686 | 1,660,328 | 2,062,964 |
| **German Giant Spitz** | 1,413,425 | 1,669,797 | 1,651,373 | 2,069,294 |
| **Pomeranian** | 1,423,291 | 1,665,841 | 1,647,144 | 2,078,092 |
| **German Spitz Mittel** | 1,451,413 | 1,672,110 | 1,652,866 | 2,073,014 |
| **Gotland Hound** | 1,378,156 | 1,672,947 | 1,655,231 | 2,097,996 |
| **German Wirehaired Pointer** | 1,423,240 | 1,660,779 | 1,642,318 | 2,098,078 |
| **Hallefors Elkhound** | 1,530,119 | 1,761,156 | 1,745,778 | 2,087,859 |
| **Harrier** | 1,291,903 | 1,652,683 | 1,640,224 | 2,097,566 |
| **Hortaya Borzaya** | 1,494,745 | 1,702,707 | 1,685,484 | 2,085,801 |
| **Hokkaido** | 1,866,895 | 2,023,324 | 2,024,277 | 2,223,790 |
| **Hovawart** | 1,524,408 | 1,692,795 | 1,673,056 | 2,110,069 |
| **Vizsla** | 1,539,721 | 1,662,008 | 1,641,340 | 2,088,355 |
| **Hamiltonstovare** | 1,378,156 | 1,672,445 | 1,656,047 | 2,093,894 |
| **Hanoverian Scenthound** | 1,542,663 | 1,705,383 | 1,683,132 | 2,111,093 |
| **Ibizan Hound** | 1,571,542 | 1,691,266 | 1,672,811 | 2,100,482 |
| **Icelandic Sheepdog** | 1,631,391 | 1,712,098 | 1,695,364 | 2,082,349 |
| **Irish Terrier** | 1,613,690 | 1,720,633 | 1,698,241 | 2,132,831 |
| **Irish Setter** | 1,425,994 | 1,657,644 | 1,636,837 | 2,094,200 |
| **Irish Water Spaniel** | 1,585,794 | 1,687,830 | 1,662,828 | 2,121,197 |
| **Japanese Spitz** | 1,608,360 | 1,721,509 | 1,703,278 | 2,090,128 |
| **Kai Ken** | 1,816,170 | 1,884,044 | 1,881,268 | 2,103,943 |
| **Kangal** | 1,625,814 | 1,736,301 | 1,723,035 | 2,086,815 |
| **Kars** | 1,687,702 | 1,772,245 | 1,757,525 | 2,100,607 |
| **Keeshond** | 1,572,698 | 1,678,971 | 1,659,869 | 2,088,814 |
| **Kerry Blue Terrier** | 1,556,894 | 1,667,452 | 1,644,594 | 2,097,249 |
| **Kintamani** | 1,727,275 | 2,068,325 | 2,072,517 | 2,249,877 |
| **Kishu** | 1,829,849 | 1,992,600 | 1,992,156 | 2,193,852 |

| | | | | |
|---|---|---|---|---|
| **Small Munsterlander** | 1,532,663 | 1,660,068 | 1,637,077 | 2,088,198 |
| **Kromfohrlander** | 1,620,825 | 1,707,955 | 1,686,108 | 2,118,113 |
| **Kuvasz** | 1,607,060 | 1,688,113 | 1,669,445 | 2,088,193 |
| **Labrador Retriever** | 1,566,636 | 1,671,412 | 1,646,749 | 2,086,764 |
| **Belgian Laekenois** | 1,527,649 | 1,691,050 | 1,671,063 | 2,095,821 |
| **Lagotto Romagnolo** | 1,559,425 | 1,655,095 | 1,633,276 | 2,076,306 |
| **Lakeland Terrier** | 1,269,032 | 1,707,015 | 1,687,829 | 2,125,381 |
| **Lancashire Heeler** | 1,559,343 | 1,676,378 | 1,654,672 | 2,092,293 |
| **Landseer** | 1,423,144 | 1,674,803 | 1,653,985 | 2,091,396 |
| **Leonberger** | 1,576,641 | 1,676,365 | 1,655,116 | 2,090,782 |
| **Catahoula Leopard Dog** | 1,525,611 | 1,639,220 | 1,618,351 | 2,077,539 |
| **Lhasa Apso** | 1,585,567 | 1,784,035 | 1,775,410 | 2,045,150 |
| **Large Munsterlander** | 1,532,663 | 1,665,304 | 1,645,623 | 2,082,536 |
| **Lowchen** | 1,599,204 | 1,691,272 | 1,669,881 | 2,104,864 |
| **Magyar Agar** | 1,522,177 | 1,665,023 | 1,643,218 | 2,090,439 |
| **Maltese** | 1,495,989 | 1,699,611 | 1,679,408 | 2,098,995 |
| **Manchester Terrier (Standard)** | 1,444,568 | 1,718,598 | 1,696,591 | 2,132,500 |
| **Maremma Sheepdog** | 1,582,656 | 1,676,406 | 1,656,712 | 2,088,890 |
| **Mastiff** | 1,404,997 | 1,686,695 | 1,668,807 | 2,113,316 |
| **Miniature Bull Terrier** | 785,014 | 1,722,512 | 1,708,715 | 2,145,531 |
| **Mountain Cur** | 1,477,764 | 1,619,579 | 1,597,949 | 2,044,039 |
| **Majorca Mastiff** | 1,296,896 | 1,665,191 | 1,647,296 | 2,094,830 |
| **Miniature Pinscher** | 1,511,804 | 1,662,179 | 1,640,443 | 2,093,378 |
| **Miniature Poodle** | 1,278,200 | 1,650,741 | 1,631,300 | 2,079,233 |
| **Miniature Schnauzer** | 1,635,407 | 1,721,309 | 1,699,108 | 2,124,985 |
| **Mudi** | 1,592,332 | 1,675,798 | 1,656,794 | 2,092,075 |
| **Norwegian Buhund** | 1,635,933 | 1,715,257 | 1,694,997 | 2,088,170 |
| **Neapolitan Mastiff** | 1,441,461 | 1,694,136 | 1,672,394 | 2,112,599 |
| **Norwegian Elkhound** | 1,530,119 | 1,733,323 | 1,715,718 | 2,103,765 |
| **Newfoundland** | 1,423,144 | 1,679,422 | 1,656,727 | 2,091,879 |
| **Norwegian Lundehund** | 1,818,217 | 1,863,881 | 1,841,361 | 2,235,905 |
| **Norrbottenspitz** | 1,601,151 | 1,697,974 | 1,680,758 | 2,065,709 |
| **Norfolk Terrier** | 1,393,679 | 1,745,182 | 1,723,342 | 2,151,228 |
| **Norwich Terrier** | 1,393,679 | 1,712,696 | 1,691,164 | 2,131,242 |
| **Nova Scotia Duck Tolling Retriever** | 1,563,289 | 1,671,223 | 1,647,518 | 2,085,208 |
| **Olde English Bulldogge** | 1,215,573 | 1,692,082 | 1,679,381 | 2,124,896 |
| **Old English Sheepdog** | 1,583,402 | 1,677,102 | 1,654,690 | 2,093,486 |
| **Old German Shepherd** | 1,381,222 | 1,648,249 | 1,632,639 | 2,065,642 |
| **Otterhound** | 1,448,203 | 1,705,021 | 1,690,389 | 2,120,067 |
| **Continental Toy Spaniel (Papillon)** | 1,307,814 | 1,658,100 | 1,637,684 | 2,081,488 |
| **Parson Russell Terrier** | 1,457,895 | 1,624,551 | 1,603,084 | 2,066,260 |
| **Pont-Audemer spaniel** | 1,421,729 | 1,648,798 | 1,627,619 | 2,081,478 |
| **Patterdale Terrier** | 1,543,034 | 1,683,189 | 1,660,651 | 2,103,058 |
| **Petit Bleu de Gascogne** | 1,413,871 | 1,643,205 | 1,624,886 | 2,078,415 |
| **Brussels Griffon (smooth)** | 1,115,238 | 1,681,683 | 1,660,923 | 2,098,573 |
| **Petit Basset Griffon Vendeen** | 1,318,161 | 1,667,989 | 1,650,090 | 2,099,949 |
| **Pekingese** | 1,572,450 | 1,775,389 | 1,764,020 | 2,066,654 |

| | | | | |
|---|---|---|---|---|
| **Pembroke Welsh Corgi** | 1,525,609 | 1,703,775 | 1,680,511 | 2,110,034 |
| **Peruvian Hairless** | 1,562,874 | 1,651,664 | 1,630,959 | 2,069,463 |
| **Continental Toy Spaniel (Phalene)** | 1,307,814 | 1,664,073 | 1,641,792 | 2,079,370 |
| **Pharaoh Hound** | 1,511,162 | 1,709,484 | 1,689,789 | 2,117,373 |
| **Picardy Shepherd** | 1,511,083 | 1,729,170 | 1,711,933 | 2,141,394 |
| **Picardy Spaniel** | 1,279,443 | 1,661,839 | 1,642,867 | 2,102,366 |
| **Plott Hound** | 1,480,686 | 1,636,366 | 1,616,749 | 2,075,106 |
| **Polish Greyhound** | 1,494,745 | 1,684,614 | 1,667,422 | 2,087,993 |
| **Polish Lowland Sheepdog** | 1,599,008 | 1,695,675 | 1,675,118 | 2,096,807 |
| **Polish Tatra Sheepdog** | 1,591,328 | 1,685,690 | 1,665,023 | 2,087,732 |
| **Poodle** | 1,278,200 | 1,651,487 | 1,632,047 | 2,070,663 |
| **Portuguese Pointer** | 1,539,140 | 1,673,605 | 1,654,056 | 2,091,380 |
| **Porcelaine** | 1,557,543 | 1,700,997 | 1,683,246 | 2,121,035 |
| **Portuguese Sheepdog** | 1,584,479 | 1,667,301 | 1,646,863 | 2,088,894 |
| **Posavac Hound** | 1,542,432 | 1,652,359 | 1,633,425 | 2,071,286 |
| **Portuguese Podengo** | 1,581,757 | 1,676,794 | 1,657,169 | 2,082,879 |
| **Portuguese Podengo Pequeno** | 1,556,928 | 1,652,278 | 1,630,863 | 2,081,152 |
| **Prague Ratter** | 1,502,220 | 1,670,970 | 1,650,772 | 2,081,456 |
| **Portuguese Water Dog** | 1,579,561 | 1,666,218 | 1,644,617 | 2,088,644 |
| **Pudelpointer** | 1,520,496 | 1,660,412 | 1,640,347 | 2,085,674 |
| **Pug** | 1,624,783 | 1,736,267 | 1,716,245 | 2,130,195 |
| **Puli** | 1,581,448 | 1,669,799 | 1,649,296 | 2,074,083 |
| **Pumi** | 1,573,416 | 1,665,255 | 1,647,025 | 2,074,494 |
| **Pyrenean Mastiff** | 1,527,484 | 1,670,683 | 1,652,226 | 2,081,918 |
| **Pyrenean Shepherd** | 1,599,617 | 1,681,198 | 1,660,091 | 2,093,618 |
| **Ratonero Bodeguero Andaluz** | 1,542,692 | 1,649,237 | 1,628,478 | 2,079,163 |
| **Rat Terrier** | 1,518,762 | 1,655,765 | 1,635,037 | 2,075,810 |
| **Redbone Coonhound** | 1,478,193 | 1,636,826 | 1,617,814 | 2,081,288 |
| **Rhodesian Ridgeback** | 1,584,309 | 1,678,630 | 1,655,422 | 2,095,075 |
| **Teddy Roosevelt Terrier** | 1,504,510 | 1,628,367 | 1,608,228 | 2,058,300 |
| **Russian Toy** | 1,560,909 | 1,655,854 | 1,636,595 | 2,075,731 |
| **Russian Hound** | 1,575,133 | 1,691,753 | 1,673,687 | 2,086,874 |
| **Irish Red and White Setter** | 1,425,994 | 1,674,764 | 1,653,807 | 2,106,237 |
| **Saarloos Wolfdog** | 1,675,219 | 1,966,339 | 1,959,437 | 2,262,979 |
| **Saluki** | 1,655,336 | 1,794,958 | 1,781,198 | 2,123,990 |
| **Samoyed** | 1,729,423 | 1,816,479 | 1,806,191 | 2,085,308 |
| **Sarplaninac** | 1,612,233 | 1,703,329 | 1,687,463 | 2,082,315 |
| **Schapendoes** | 1,602,070 | 1,689,973 | 1,667,240 | 2,107,102 |
| **Sealyham Terrier** | 1,532,756 | 1,701,094 | 1,679,059 | 2,115,029 |
| **Segugio Italiano** | 1,521,347 | 1,647,400 | 1,626,909 | 2,083,792 |
| **Chinese Shar-Pei** | 1,803,596 | 1,975,712 | 1,975,357 | 2,167,771 |
| **Shiba Inu** | 1,816,559 | 1,980,849 | 1,980,924 | 2,193,241 |
| **Shih Tzu** | 1,572,450 | 1,756,460 | 1,745,166 | 2,046,157 |
| **Shiloh Shepherd** | 1,163,733 | 1,713,838 | 1,705,384 | 2,121,896 |
| **Shikoku** | 1,842,164 | 2,090,039 | 2,092,595 | 2,286,807 |
| **Skye Terrier** | 1,648,946 | 1,726,252 | 1,702,262 | 2,134,834 |
| **Silken Windhound** | 1,494,490 | 1,692,574 | 1,671,881 | 2,105,851 |

| | | | | |
|---|---|---|---|---|
| **Swedish Lapphund** | 1,626,936 | 1,716,146 | 1,696,914 | 2,085,392 |
| **Slovak Cuvac** | 1,599,025 | 1,694,080 | 1,674,826 | 2,097,183 |
| **Slovensky Kopov** | 1,597,496 | 1,682,945 | 1,660,981 | 2,100,998 |
| **Sloughi** | 1,574,476 | 1,690,072 | 1,672,591 | 2,081,070 |
| **Slovakian Wirehaired Pointer** | 1,458,887 | 1,666,708 | 1,646,282 | 2,091,535 |
| **Saint Miguel Cattle Dog** | 1,558,803 | 1,656,341 | 1,635,691 | 2,069,036 |
| **Small Swiss Hound** | 1,559,471 | 1,699,561 | 1,677,876 | 2,114,360 |
| **Spinone Italiano** | 1,552,600 | 1,652,200 | 1,632,408 | 2,079,181 |
| **Spanish Mastiff** | 1,570,948 | 1,666,401 | 1,647,101 | 2,080,353 |
| **Spanish Water Dog** | 1,557,400 | 1,657,217 | 1,637,999 | 2,075,647 |
| **Sarabi** | 1,677,412 | 1,766,890 | 1,752,533 | 2,106,539 |
| **South Russian Ovcharka** | 1,619,668 | 1,712,152 | 1,694,139 | 2,103,369 |
| **Shetland Sheepdog** | 1,305,666 | 1,698,456 | 1,678,344 | 2,121,209 |
| **Standard Schnauzer** | 1,597,505 | 1,683,567 | 1,662,397 | 2,087,853 |
| **Staffordshire Bull Terrier** | 1,484,020 | 1,676,571 | 1,659,894 | 2,110,075 |
| **Saint Bernard** | 1,576,641 | 1,681,397 | 1,659,276 | 2,093,308 |
| **Stabyhoun** | 1,627,928 | 1,707,620 | 1,683,432 | 2,118,738 |
| **Saint-Usuge Spaniel** | 1,421,729 | 1,641,834 | 1,621,149 | 2,080,620 |
| **Sussex Spaniel** | 1,632,441 | 1,748,102 | 1,723,465 | 2,163,092 |
| **Swedish Vallhund** | 1,636,875 | 1,718,077 | 1,697,261 | 2,108,122 |
| **Swedish Elkhound** | 1,512,759 | 1,699,257 | 1,679,023 | 2,089,519 |
| **Swiss Hound** | 1,458,197 | 1,670,230 | 1,650,481 | 2,091,526 |
| **Swedish White Elkhound** | 1,512,759 | 1,722,971 | 1,703,944 | 2,096,412 |
| **Tazi** | 1,655,336 | 1,769,507 | 1,756,808 | 2,103,636 |
| **Thai Ridgeback** | 1,727,275 | 2,000,901 | 1,999,847 | 2,203,478 |
| **Tibetan Spaniel** | 1,670,339 | 1,835,461 | 1,827,310 | 2,085,835 |
| **Tibetan Terrier** | 1,663,099 | 1,747,748 | 1,732,245 | 2,072,655 |
| **Tamaskan** | 1,478,816 | 1,759,795 | 1,752,611 | 2,057,238 |
| **Tosa Inu** | 1,594,706 | 1,724,364 | 1,711,283 | 2,009,837 |
| **Toy Poodle** | 1,344,958 | 1,646,908 | 1,627,052 | 2,070,613 |
| **Turkish Mastiff** | 1,625,814 | 1,734,701 | 1,719,021 | 2,091,816 |
| **Treeing Walker Coonhound** | 1,475,916 | 1,635,643 | 1,616,182 | 2,082,492 |
| **Toy Fox Terrier** | 1,513,563 | 1,648,017 | 1,624,411 | 2,085,730 |
| **Volpino Italiano** | 1,554,115 | 1,699,126 | 1,680,358 | 2,090,745 |
| **Weimaraner** | 1,458,887 | 1,693,488 | 1,671,444 | 2,110,554 |
| **Welsh Terrier** | 1,317,199 | 1,679,356 | 1,659,270 | 2,103,829 |
| **Whippet** | 1,494,490 | 1,681,951 | 1,658,850 | 2,113,472 |
| **Wirehaired Pointing Griffon** | 1,541,944 | 1,671,725 | 1,651,537 | 2,106,995 |
| **Welsh Springer Spaniel** | 1,580,804 | 1,693,527 | 1,670,992 | 2,112,770 |
| **Xoloitzcuintli** | 1,534,222 | 1,630,876 | 1,612,681 | 2,059,468 |
| **Yorkshire Terrier** | 1,370,539 | 1,669,889 | 1,649,305 | 2,091,414 |
| **Zagar** | 1,558,818 | 1,666,211 | 1,647,602 | 2,075,948 |
| **Turkish Zerdava** | 1,624,104 | 1,716,636 | 1,701,487 | 2,084,392 |

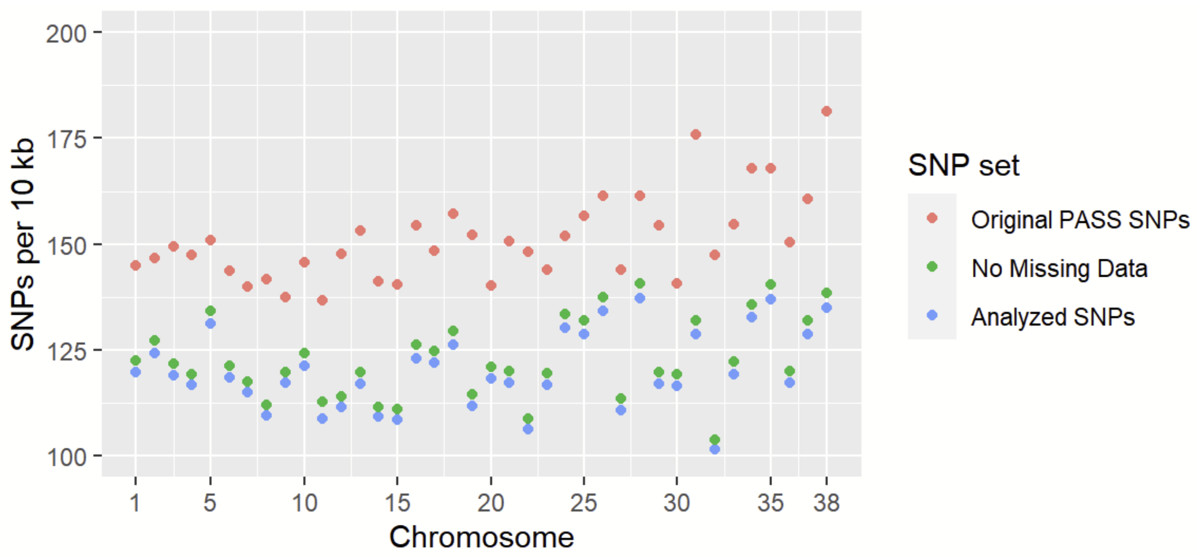


**Figure S1. Density of SNPs included in canine analysis**
The density of SNPs per 10 kb is shown for different filtration sets across the 38 canine autosomes. The red points represent all sites listed as 'PASS' in the initial VCF file. The green points represent sites with no missing data in the 1,929 considered samples. The blue points represent the final variant set used for downstream analysis.

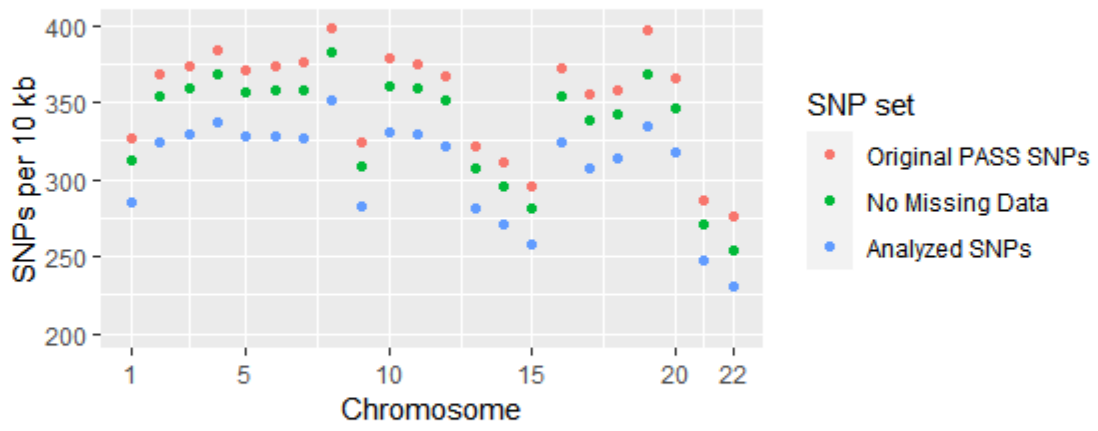


**Figure S2. Density of SNPs from the Human 1000 Genomes Project**
The rate of original PASS SNPs, SNPs with no missing data, and analyzed SNPs, from the high-coverage 1000 Genomes Project data set is shown.

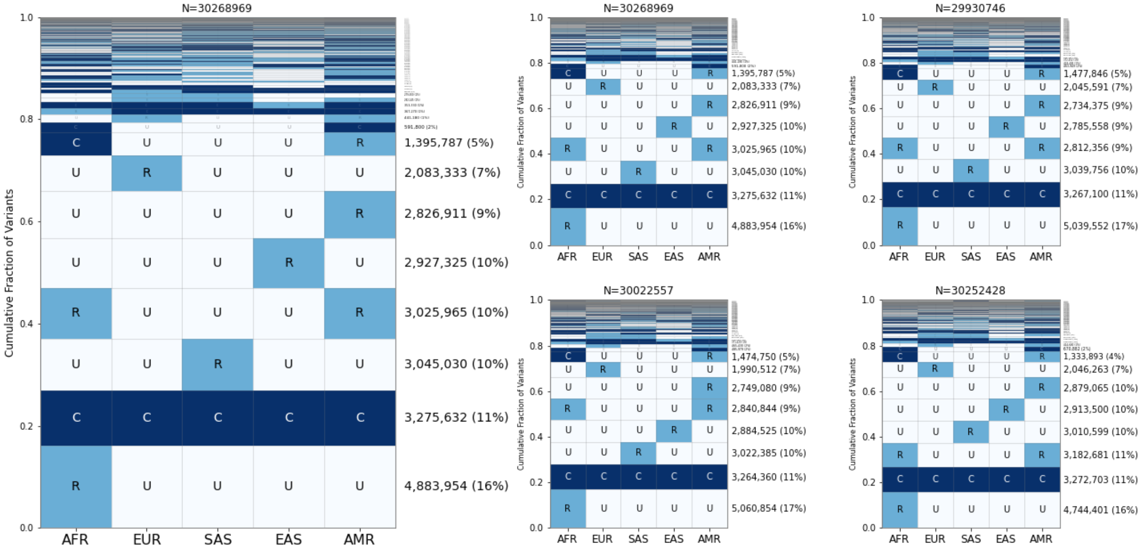


**Figure S3. Human GeoVar analysis with reduced sample sizes**

A similar analysis to that of Figure 3 but with the human 1000 Genomes Project data, arranged in the same five populations as Biddanda *et. al*. Population codes: AFR=Africa, EUR=Europe, SAS=South Asia, EAS=East Asia, AMR=Americas. A random selection of 50 individuals per group was analyzed.

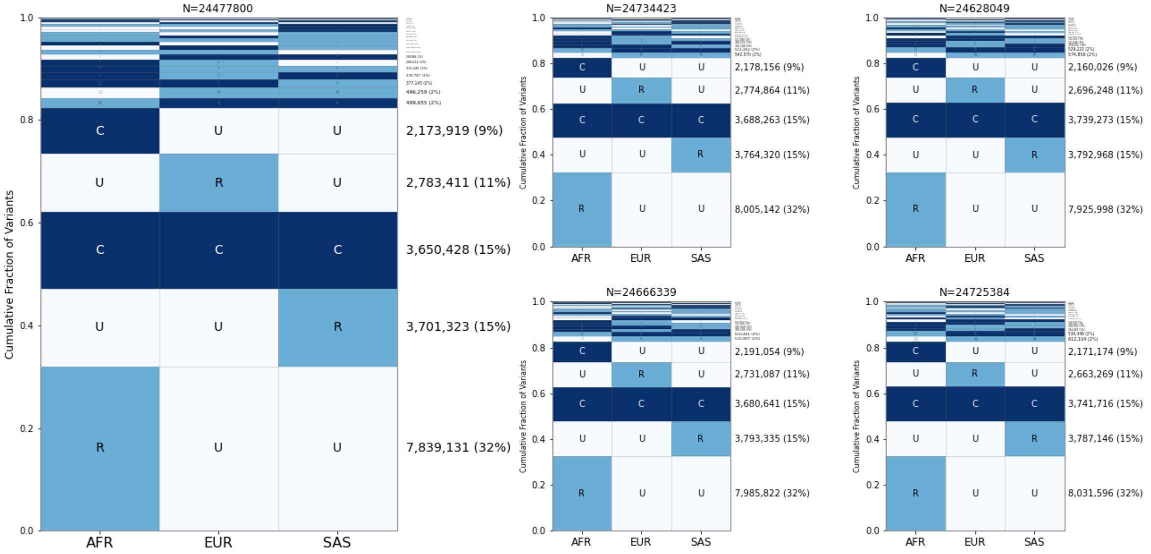


**Figure S4. Human GeoVar analysis with three groups**

The same analysis as Figure S2, but with three populations instead of five. The South and East Asian populations have been combined into ‘SAS’, and the America population has been removed.

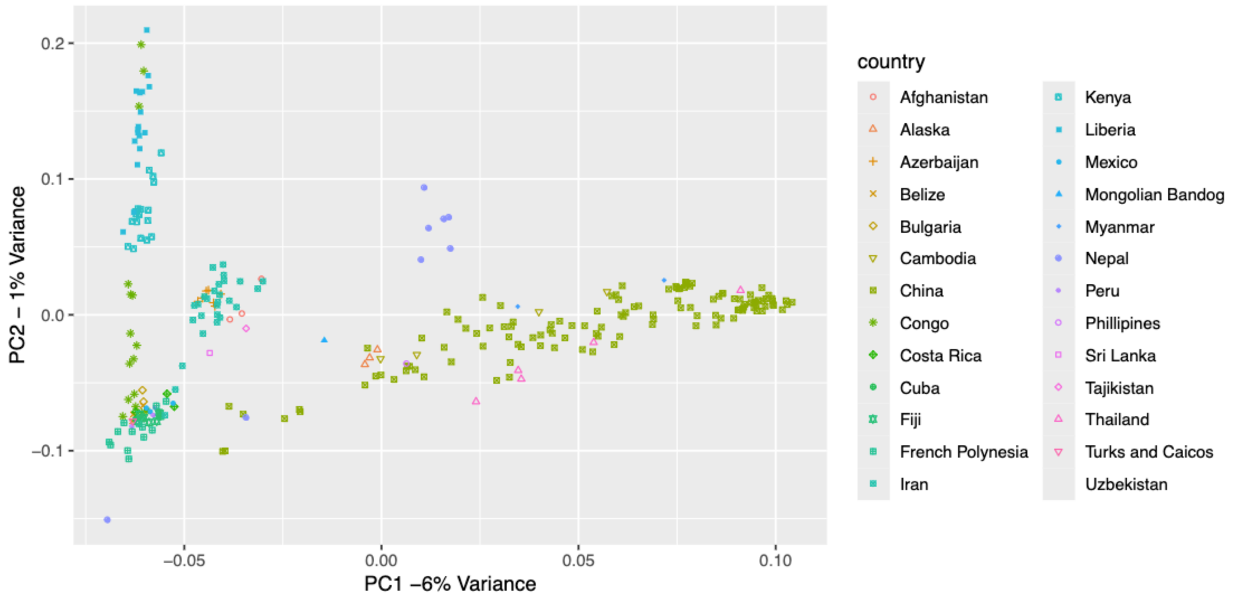


**Figure S5. Principal component analysis of village dogs**

The three groups analyzed as populations are shown with the points colored based on sample location. Apart from the Chinese village dogs, which show a wide range of PC1 values, and were categorized according to the grouping shown, all village dogs from one country were categorized into one of the three categories, or not categorized at all. The categories are: Africa = Congo, Liberia, Kenya; Central Asia = Afghanistan, Azerbaijan, Bulgaria, Iran, Tajikistan, Uzbekistan, East Asia =  Cambodia, China (most samples), Myanmar, Nepal, Philippines, Sri Lanka, Thailand. The exact samples in each category are listed in Table S1.
